# N- and T-genes framework for pathway deregulation: Taxonomy, progression, and protection in cancer

**DOI:** 10.64898/2026.11.13.766402

**Authors:** Joan Nieves, Dania Vázquez, Augusto González

## Abstract

**Background:** Pathway analysis is widely used to interpret gene expression, but conventional enrichment depends on arbitrary thresholds and does not quantify deregulation magnitude. Recent reformulations based on average log-fold changes remove the threshold but still average over all annotated genes, diluting the signal from the genes that actually drive the normal-to-tumor transition. The N/T framework, which identifies genes with exclusive expression intervals in normal (N) or tumor (T) samples, offers a selective alternative, but it had not been extended to pathway-level quantification, taxonomy, or cross-cancer comparison.

**Methods:** We define effective activation frequencies for Reactome pathways based on N- and T-genes: for each sample and pathway, the effective N (T) frequency is the fraction of annotated N (T) genes active in a normal (tumor) sample. The analysis proceeds in three explicit steps: (i) cohort averages identify the pathways important in each tissue; (ii) averages over hierarchical-clustering groups, ordered by the number of active genes, reveal sequences of pathway activation and deactivation along progression; (iii) a minimum spanning tree (MST) built on pathway-frequency vectors—independent of the number of active genes—defines the molecular taxonomy, and branch averages identify subtype-specific programs. We apply the framework to the full set of Reactome pathways in prostate adenocarcinoma (PRAD), lung adenocarcinoma (LUAD), liver hepatocellular carcinoma (LIHC), and glioblastoma (GBM), analyzing both tumor and normal compartments.

**Results:** In PRAD tumors (n=499), hierarchical clustering identifies four ordered progression groups and the MST resolves six main molecular subtypes comprising 406 tumors (81.4%): B1 (42.1%; MET/NFE2L2/energy metabolism), B2 (13.8%; RAS/apoptosis/telomere), and four smaller subtypes defined by PTEN (B3), DNA repair and lipid metabolism (B4), inflammation and LXR-mediated lipid metabolism (B5), and neuronal development and base excision repair (B6). An eight-gene perfect T panel reproduces the MST subtypes at gene level. Analysis of normal prostatic tissue (n=52) reveals a hierarchy of protection: immune response, apoptosis, and senescence are lost or partially lost early, DNA repair at an intermediate stage, and cell cycle regulation and receptor tyrosine kinase signaling last; an eleven-gene N panel reproduces this hierarchy at gene level. The cross-cancer comparison shows that this architecture is a general property: DNA repair, cell cycle, and RTK protection are maintained in normal tissue across PRAD, LUAD, LIHC, and GBM, while the specific pathways lost in normals and gained in tumors are tissue-specific (MET/NFE2L2 in PRAD; NTRK/LXR/phagosome in LIHC; surfactant/FGFR/glycosylation in LUAD; TP53 loss/lipid metabolism/glycosylation in GBM). Taxonomic heterogeneity decreases from PRAD (six subtypes) to LIHC (two) to LUAD and GBM (one dominant program), paralleling an increase in the number of activated T-genes.

**Conclusions:** The framework eliminates dilution by non-informative genes, quantifies pathway deregulation sample by sample, and unifies tumor taxonomy with a molecular indicator of progression. Its most distinctive contribution is the joint analysis of normal and tumor compartments across cancer types, which links loss of protection in normal tissue to gain of progression programs in tumors and reveals both shared and tissue-specific cancer mechanisms. The method is general and applicable across cancer types, although quantitative results depend on the underlying cohort.

## 1. Introduction

### 1.1 Background

Pathway analysis has become a cornerstone of systems biology for interpreting gene expression data [1]. The conventional approach relies on hypothesis-testing schemes, typically enrichment analysis, where a set of differentially expressed genes is first selected and then evaluated for over-representation in a given pathway. This approach has two well-known limitations. First, the result depends on the threshold and size of the selected gene set, which introduces an element of arbitrariness. Second, enrichment analysis only provides a probability of over-representation; it does not quantify the magnitude of deregulation. A pathway may be significantly enriched but only mildly deregulated, or vice versa.

Recently, Costa, Nieves, and Gonzalez [2] proposed a reformulation of pathway analysis based on the average log-fold change of the genes comprising each pathway. This method removes dependence on the selected gene set, allows direct quantification of deregulation magnitude, and enables sample-by-sample analysis, which in turn facilitates sample classification. However, this approach still averages over all genes in the pathway, including those that do not participate in the biological transition of interest. If a pathway contains twenty annotated genes but only two of them are actually involved in the transition from normal tissue to tumor, the remaining eighteen genes dilute the signal and obscure the pathway’s true contribution.

In parallel, Gil et al [3.4] introduced the concept of N- and T-genes. These are genes whose expression exhibits exclusive intervals in normal (N) or tumor (T) samples, respectively. The identification relies on statistically significant exclusive expression intervals, and expression is discretized into three states: e = −1 for N-active, e = +1 for T-active, and e = 0 for the common state. A gene may have a normal-exclusive interval, a tumor-exclusive interval, or both. In the latter case, the gene belongs simultaneously to the N and T categories, and is treated as such throughout this work. This framework has allowed the decomposition of carcinogenesis into two distinct phases: somatic evolution, characterized by the loss of N-gene activity, and tumor progression, characterized by the gain of T-gene activity. The number of active N-genes in a normal sample or active T-genes in a tumor sample serves as an indicator of the respective phase.

The convergence of these two lines—the need to avoid dilution and the availability of genes with exclusive intervals—defines the starting point of this work: if only N- and T-genes carry information about the normal-to-tumor transition, then the natural measure of a pathway’s activity is not an average over all its annotated genes, but an **effective frequency** restricted to the N- and T-genes it contains. Such a frequency would be, by construction, a normalized measure, directly comparable across pathways and samples, and separable into two complementary readings: the presence of protection in normal tissue and the gain of progression in tumor tissue.

### 1.2 Pathway-based taxonomy: prior work and remaining limitations

The use of pathway-level information for tumor classification is not new. Bild et al. [5] demonstrated that gene expression signatures reflecting oncogenic pathway activation status can define tumor subtypes and predict therapeutic response. Gatza et al. [6] built a pathway-based classification of breast cancer, identifying subgroups that shared patterns of pathway activity and showed distinct clinical and biological properties. Drier et al. [7] introduced Pathifier, an algorithm that computes a Pathway Deregulation Score (PDS) per sample and pathway. They showed that clustering on PDS identifies clinically relevant subtypes in glioblastoma and colon cancer, including subgroups not captured by standard expression-based methods. Livshits et al. [8] applied Pathifier to the METABRIC breast cancer dataset, stratifying tumors into nine types with distinct PDS profiles.

However, these approaches share a common limitation: they compute pathway-level scores using all annotated genes in the pathway. Bild et al. and Gatza et al. use trained classifiers, where genes receive empirical weights derived from training data. Costa et al. use a uniform average of log-fold changes, where all genes weigh equally. Pathifier uses a geometric distance along a principal curve, where the weight of each gene is implicit in the curve construction. In all cases, a pathway may be scored as moderately deregulated simply because many of its genes show small changes, even if a few key genes are strongly and exclusively involved in the transition. This dilution effect is precisely what the N/T framework is designed to overcome, by restricting the analysis to genes with exclusive expression intervals, which are the ones actually driving the transition.

To this methodological limitation is added a limitation of scope. Pathway-based taxonomies developed so far—including the earlier ones—have been formulated and validated within a single tumor type and have focused exclusively on the tumor compartment. They did not analyze normal tissue, and therefore did not address the relationship between loss of protection in normal samples and gain of progression programs in tumors.

### 1.3 Motivation

Motivated by the above, we propose a reformulation of pathway analysis that is applicable homogeneously across several cancer types and that analyzes both the normal and the tumor compartments. It restricts the analysis to N- and T-genes annotated in pathways, and defines an effective activation frequency per pathway. This reformulation has several conceptual advantages. It eliminates dilution by non-informative genes, it provides a direct and normalized measure of pathway deregulation, and it allows separate evaluation of loss of protection in normal samples and gain of progression in tumor samples. Moreover, it connects naturally with the N/T framework and its decomposition of carcinogenesis into two phases.

Beyond the methodological reformulation, this work addresses a broader conceptual problem in cancer genomics: the relationship between classification (taxonomy) and temporal evolution (staging). Traditionally, tumor groups are defined by a set of mutational drivers, while staging comes from clinical pathology. Here we propose a fully molecular alternative: pathways are used to classify samples, while the number of activated genes is used as an indicator of temporal evolution. This unifies taxonomy and staging into a single molecular framework, although we emphasize that the number of activated genes is an indicator, not a clinical staging tool.

### 1.4 The three-step analytical logic

A distinctive feature of this work is that the analysis is organized in three explicit, sequential steps, each answering a different biological question with a different level of aggregation:

1. **Global averages (whole cohort).** The average effective frequency of each pathway over the complete cohort (all tumors, or all normals) identifies the pathways that are important in that tissue: the programs most frequently deregulated in tumors, or most robustly maintained as protection in normals.
2. **Averages over hierarchical clustering groups.** The samples are ordered by the number of active genes (T-genes in tumors, N-genes in normals), and clustered on their pathway frequency vectors. Averaging each pathway over the resulting groups, and comparing consecutive groups along the progression order, reveals the sequences of pathway activation and deactivation: which programs appear early, which appear late, which disappear, and which show transient peaks.
3. **MST taxonomy and branch averages.** The minimal spanning tree, built on the same pathway frequency vectors but without using the number of active genes in its construction, provides the taxonomy: its main branches define molecular subtypes, and averaging each pathway over the samples of a branch identifies the pathways characteristic of that subtype.

This ordering—global, then clustering-based sequences, then MST-based taxonomy—is used consistently throughout the paper and its Supplementary Material.

### 1.5 Objectives

The objectives of this study follow directly from the three-step analytical logic described above and from the N/T framework. They are stated in the order in which they are addressed in the paper.

First, to formally define the effective activation frequency of a pathway in a sample, based on the N- and T-genes annotated in Reactome. This definition is the methodological core of the paper: it replaces the average over all annotated genes with a frequency restricted to the genes that actually carry information about the normal-to-tumor transition.

Second, to compute average effective frequencies per pathway in normal and tumor cohorts across four cancer types—PRAD, LUAD, LIHC, and GBM—using PRAD as the detailed case study and the other three tissues as the basis for the cross-cancer comparison. The four tissues differ in cohort size and in the availability of normal samples, and these differences are stated explicitly in the analyses that follow.

Third, to order the samples along the number of activated genes as an indicator of temporal evolution. The groups obtained in this step describe progression.

Fourth, to build a molecular taxonomy with the MST on pathway frequency vectors, and to characterize the resulting branches by their pathway deregulation profiles. The branches are then compared with previous pathway-based taxonomies.

Fifth, to characterize the normal compartment through global averages of protection programs and, where the data allow, through the sequence of their loss along progression. The hierarchy of protection is supported primarily by the global averages of Step 1; the group-level sequence is reported as a qualitative, complementary view, because the trajectories of several protection programs in normal tissue are not monotonic.

Sixth, to compare protection mechanisms in normal tissue and progression programs in tumors across the four cancer types, identifying shared and tissue-specific features. This comparison is the most distinctive contribution of the work, because previous pathway-based taxonomies were developed and validated within a single tumor type.

Seventh, to verify the pathway-level results against the complete N- and T-gene panels defined in Gil et al. [4], connecting the pathway-level and gene-level views of the taxonomy and of the protection hierarchy.

## 2. Methods

### 2.1 Data

RNA-seq gene expression data in FPKM format are obtained from The Cancer Genome Atlas (TCGA) public repository. We focus on three cancer types for the taxonomy and staging analysis: prostate adenocarcinoma (PRAD) [9], lung adenocarcinoma (LUAD) [10], and liver hepatocellular carcinoma (LIHC) [11]. These three cohorts contain sufficient numbers of both normal and tumor samples to identify N- and T-genes and to perform a robust classification analysis. Glioblastoma (GBM) [12] is qualitatively analyzed, because the TCGA GBM cohort contains very few normal samples, which limits the identification of N-genes but does not prevent the analysis of T-genes. The main text presents PRAD as the detailed case study and the cross-cancer comparison across the four tissues. Extended per-tissue analyses for LIHC, LUAD, and GBM are provided in the Supplementary Material.

Pathway annotations are taken from Reactome [13], version v91 or higher. We include the full set of pathways in the Reactome hierarchy. This provides a finer-grained description of the biological programs active in each sample, and it allows the taxonomy and staging analysis to capture subpathway-level deregulation events that would be invisible at the top level.

### 2.2 Identification of N- and T-genes

The identification of N- and T-genes follows the method described in Gil et al. [3,4]. For each gene, expression distributions in normal and tumor samples are compared. Exclusive intervals that are significantly populated are identified using Fisher’s exact test. The significance thresholds are set at 5% for N-genes and 10% for T-genes. A gene is classified as an N-gene if it has a normal-exclusive interval, and as a T-gene if it has a tumor-exclusive interval. A gene may belong to both categories simultaneously, in which case it is treated as an N-gene when analyzing normal samples and as a T-gene when analyzing tumor samples. Genes with no exclusive intervals are not used in this analysis.

### 2.3 Filtering by Reactome annotation

Only N- and T-genes that are annotated in Reactome are retained for the pathway analysis. For each pathway P, we record the number of N-genes, denoted nN(P), and the number of T-genes, denoted nT(P). A gene that belongs to both categories is counted in both nN(P) and nT(P). Pathways with no N- or T-genes are excluded from the analysis, or reported with frequency zero if they are of particular interest.

### 2.4 Definition of effective frequencies

For a given sample s and a pathway P, we define the effective N frequency as the number of N-genes that are active in sample s, provided the sample corresponds to a normal tissue, divided by nN(P). Similarly, the effective T frequency is defined as the number of T-genes that are active in sample s, a tumor sample, divided by nT(P). Here, active means that the discretized expression state is e = −1 for N-genes and e = +1 for T-genes.

In normal samples, the N frequency measures the presence of protection mechanisms. In tumor samples, on the other hand, the T frequency measures the gain of progression mechanisms. The two analyses are performed separately and independently, reflecting the two distinct phases of carcinogenesis.

### 2.5 Step 1 — Cohort averages and identification of important pathways

For each pathway, we compute the average effective N frequency over all normal samples and the average effective T frequency over all tumor samples. These averages are computed separately for each cancer type, allowing cross-cancer comparisons. The average pathway frequencies in the sample population are then analyzed to extract the most important (most frequently deregulated) pathways in each tumor type. These pathways represent the dominant biological programs active in the cohort. In the same way, the most important (most frequently regulated) pathways in normal tissue represent the main mechanisms of protection against cancer.

This first step answers the question: *which biological programs dominate in this tissue?* It is reported as a table of representative pathways grouped by general mechanism, and it constitutes the reference against which the group-level and branch-level averages of the following steps are interpreted.

### 2.6 Step 2 — Hierarchical clustering and sequences of activation/deactivation

Using the vectors of pathway frequencies in individual samples, we perform a hierarchical clustering analysis. Samples are ordered by the number of active genes (T-genes in tumors, N-genes in normals), which serves as a temporal indicator, and the resulting groups are interpreted as stages of progression.

The key output of this step is not the partition itself, but the **sequences** it reveals. For every pathway, we compare its average frequency across the ordered groups and classify its trajectory using an explicit relative criterion:

- **Active in a group:** the group-average frequency exceeds the global cohort average for that pathway.
- **Appreciable change:** the absolute difference between consecutive groups is at least 0.10.
- **Trajectory classes:** *gains* (monotonic or near-monotonic increase with at least one appreciable step), *loses* (monotonic or near-monotonic decrease with at least one appreciable step), *transient peak* (maximum in an intermediate group with an appreciable fall on both sides), and *stable* (no appreciable change).
- **Onset:** the first group, in progression order, in which the pathway exceeds its global average.

In normal samples, the same criterion is applied with inverted semantics: a high frequency indicates *maintained protection*, and a frequency that falls along the progression order indicates *lost protection*.

This second step answers the question: *in what order are programs gained or lost?* It is reported as a table of representative pathways per mechanism, with their frequencies in each ordered group, their trajectory class, and their onset.

### 2.7 Step 3 — MST taxonomy and branch averages

A key conceptual point of this work is that classification and temporal evolution are intertwined, but they must not be conflated. The MST algorithm is used precisely to avoid the trivial connection between groups and progression. If samples were clustered by their number of activated genes, the result would be a progression classification, not a taxonomy. The MST instead clusters samples by their pathway frequency vectors, which are independent of the number of activated genes. The fact that the resulting branches correlate with progression is a validation of the method, not a limitation.

The MST algorithm works only with the distance between pairs of samples, which in our case is determined by the frequency vectors. It was originally formulated by Boruvka [14] and later rediscovered by Jarnik [15], Kruskal [16], and Prim [17]. Its application to gene expression clustering was pioneered by Xu et al. [18], who showed that MST-based methods can identify meaningful partitions in microarray data. Subsequent work extended this approach to iterative class discovery and feature selection [19] and to marker gene selection for sample classification [20]. More recently, Gagolewski et al. [21] provided a comprehensive review showing that MST-based clustering methods often outperform K-means, Gaussian mixtures, spectral clustering, and hierarchical agglomerative procedures. In the biomedical context, Kolles et al. [22] applied MST to glioma grading, demonstrating that MST parameters can quantify histological features such as cellularity and nuclear crowding. Their work shows that MST topology can encode information about a progressive process without using the progression variable in the construction of the tree. Our method extends this idea to the molecular domain, using pathway frequency vectors as the distance metric and the number of activated genes as the temporal indicator. Zou [23] introduced perturbed MST constructions for single-cell trajectory analysis, which is relevant because our approach also deals with continuous transitions.

Once the main branches are identified, we determine the most important pathways in each branch by comparing the average effective frequency vectors across branches (for tumors). Pathways that are significantly more frequent in a branch represent the biological programs that characterize that molecular subtype.

This third step answers the question: *what functionally defines each subtype?* It is reported as a table of representative pathways per branch, with the branch frequency, the global frequency, and the enrichment ratio (branch/global).

### 2.8 Verification with complete gene panels

We verify whether the genes of the complete panels defined in Gil et al. [4] are capable of parameterizing the main groups. The complete panels are sets of genes with 100% sensitivity and 100% specificity within the training set, and they provide a complementary, gene-level view of the taxonomy. If the panel genes are able to reproduce the main groups, this would strengthen the consistency of the framework. This analysis was left pending in [4] and is addressed here. For PRAD, the panels comprise eight T-genes and eleven N-genes; their activation is computed per clustering group and per MST branch, and compared with the pathway-level results of Steps 2 and 3.

### 2.9 Cross-cancer comparison

The cross-cancer comparison applies the three steps of Sections 2.5–2.7 independently to each cohort (PRAD, LIHC, LUAD, GBM) and then compares the resulting summaries across tissues at three levels.

For each tissue, the protection programs in normals are matched to the dominant tumor programs of the same tissue by shared mechanism labels. Correspondences are reported as qualitative associations, not as causal claims, and the direction (loss in normals vs. gain in tumors) is stated explicitly for each pair.

**Glossary. Terminology used in this work.**

| Term | Definition | Labels used |
|---|---|---|
| Branch | Connected component of the MST; graph-theoretic object | B1–B6 (PRAD tumors); branches in LIHC, LUAD, GBM |
| Subtype | Biological interpretation of a branch with a coherent pathway profile | B1–B6 when interpreted biologically |
| Group | Cluster from hierarchical clustering, ordered by number of active genes | G1–G4 (tumors); G1–G3 (normals) |
| Cluster | Generic term for the output of a clustering algorithm | — |

## 3. Results

### 3.1 Step 1 — Pathways important in PRAD tumors

The global averages over the 499 PRAD tumors identify the dominant biological programs of the cohort. Figure 1 upper panel shows the frequency of active pathways in tumors. Grouped by general mechanism, the most important pathways are: MET receptor activation (0.559) and NTF3–NTRK2 signaling (0.511) among receptor tyrosine kinase programs; NFE2L2 regulation of inflammation- and ER-stress-associated genes (0.549) among oxidative-stress programs; utilization of ketone bodies (0.501) and ketone body metabolism (0.400) among energy-metabolism programs; transport of fatty acids (0.439) and glycerophospholipid catabolism (0.355) among lipid programs; axonal growth stimulation (0.507) and CREB1 phosphorylation via CaMKII/CaMKK/CaMKIV (0.471) among neurotrophin/axonal programs; the AIM2 inflammasome (0.347) among inflammatory programs; and, at lower global frequencies, apoptotic, DNA-repair and telomere programs (SMAC– IAP, 0.127; defective mismatch repair MSH6, 0.182; alternative lengthening of telomeres, 0.136). These global values constitute the reference for the sequence and taxonomy analyses below.

**Figure 1.**
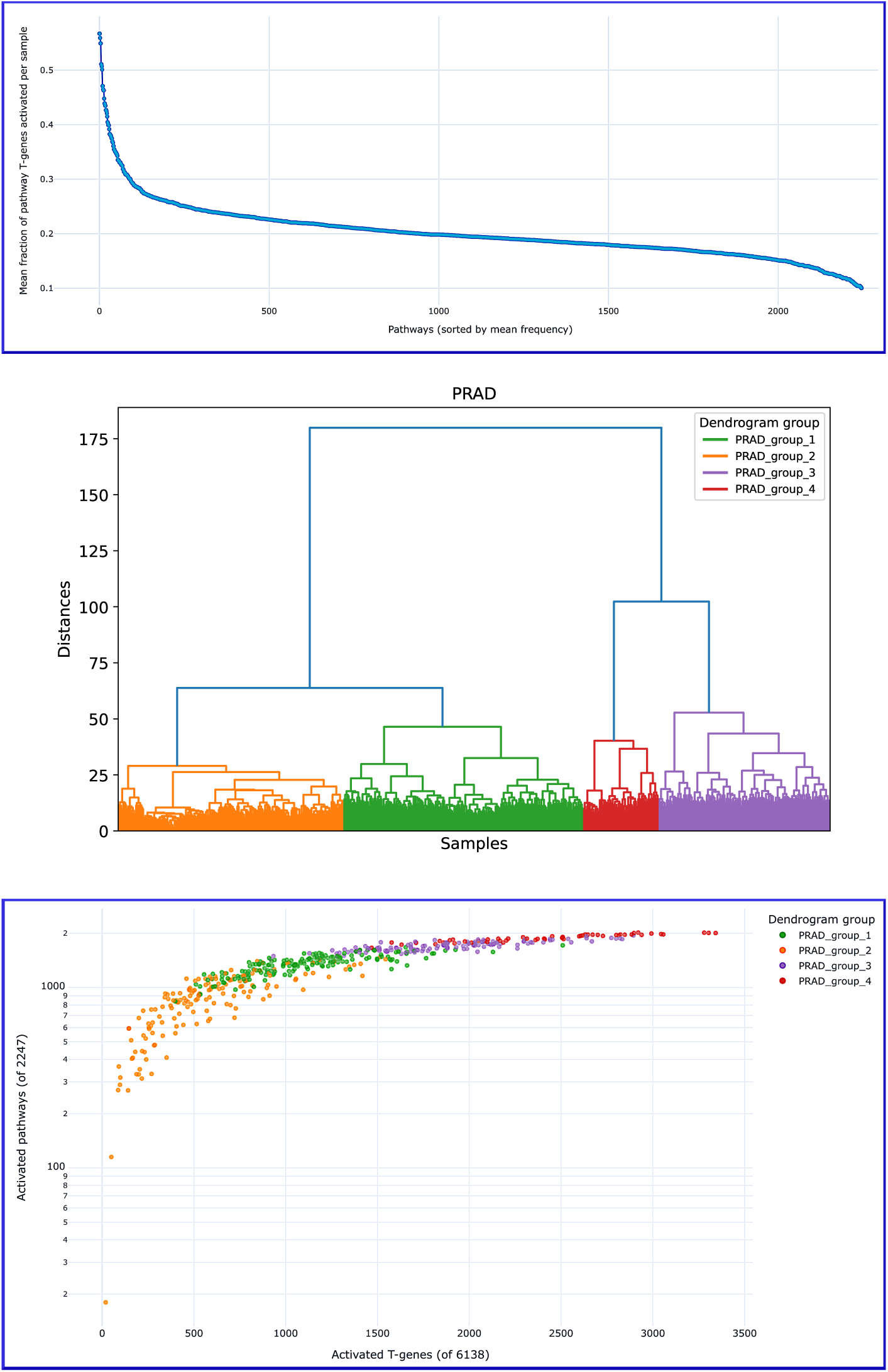
Clustering of PRAD tumor samples according to the effective T frequency vectors. Upper panel: mean frequency of pathways in PRAD tumors, sorted by mean frequency (Step 1). Middle panel: dendrogram showing four groups of samples. Lower panel: number of activated pathways as a function of the number of active T-genes, with samples colored according to the clusters. The groups, ordered G2→G1→G3→G4, represent stages in tumor progression (Step 2). The number of activated pathways saturates at high T-gene counts.

### 3.2 Step 2 — Sequences of activation/deactivation in PRAD tumors

Hierarchical clustering of the PRAD tumor samples on their effective T frequency vectors identifies four groups, as shown in the dendrogram of Figure 1, central panel. Figure 1 lower panel shows that the groups may be ordered by the number of active T-genes as G2 → G1 → G3 → G4.

Comparing pathway averages across these ordered groups reveals a staggered gain of programs (Table 1).

**Table 1.** Sequences of pathway activation/deactivation along PRAD tumor progression (G2→G1→G3→G4). For each mechanism, a representative pathway is shown with its average effective T frequency in each group, its trajectory class, and its onset (first group exceeding the global average). Criterion: active if group frequency > global.

| **Mechanism** | **Representative pathway** | **G2** | **G1** | **G3** | **G4** | **Trajectory** | **Onset** |
|---|---|---|---|---|---|---|---|
| RTK / MET | MET Receptor Activation | 0.28 | 0.67 | 0.66 | 0.83 | Gains | G1 |
| Oxidative stress | NFE2L2 regulating ER-stress genes | 0.29 | 0.49 | 0.78 | 0.98 | Gains | G3 |
| Energy metabolism | Utilization of Ketone Bodies | 0.3 | 0.51 | 0.68 | 0.7 | Gains | G1 |
| Neurotrophins | NTF3 activates NTRK2 (TRKB) | 0.2 | 0.57 | 0.76 | 0.71 | Gains (plateau) | G1 |
| Lipid transport | Transport of fatty acids | 0.18 | 0.48 | 0.54 | 0.85 | Gains | G1/G3 |
| Axonal | Axonal growth stimulation | 0.32 | 0.46 | 0.66 | 0.85 | Gains | G3 |
| Inflammation | The AIM2 inflammasome | 0.11 | 0.49 | 0.53 | 0.44 | Transient peak (G1–G3) | G1 |
| Apoptosis | SMAC (DIABLO) binds to IAPs | 0.04 | 0.02 | 0.11 | 0.75 | Late gain | G4 |
| DNA repair | Defective Mismatch Repair MSH6 | 0.01 | 0.27 | 0.37 | 0.71 | Gains | G1 |
| Telomere | Alternative Lengthening of Telomeres | 0.03 | 0.02 | 0.09 | 0.94 | Late gain | G4 |

The sequence is biologically coherent: RTK/MET, energy metabolism, DNA repair and neurotrophin programs are gained early (G1); NFE2L2 and axonal programs in G3; apoptosis and telomere maintenance only in G4, i.e., at an advanced stage. Inflammation shows a transient peak in the intermediate groups. The saturation of activated pathways at high T-gene counts, visible when the number of activated pathways is plotted against the number of active T-genes, reflects the finite number of pathways that can be deregulated.

### 3.3 Step 3 — MST taxonomy and molecular subtypes of PRAD

The MST constructed from the effective T frequency vectors of the 499 PRAD tumors has 26 branches in total (Figure 2). The six largest branches contain 406 samples (81.4%) and define six molecular subtypes with exact sizes **B1 = 210, B2 = 69, B3 = 35, B4 = 34, B5 = 31, B6 = 27** ; the remaining 93 samples are distributed across 20 minor branches, each containing fewer than 21 samples. The root, represented as a red point, is a fictitious sample with zero activated T-genes; the tree flows mainly in the direction of increasing numbers of active T-genes, confirming that this number is a good temporal indicator. Averaging pathway frequencies over each branch identifies the programs characteristic of each subtype (Table 2).

**Figure 2.**
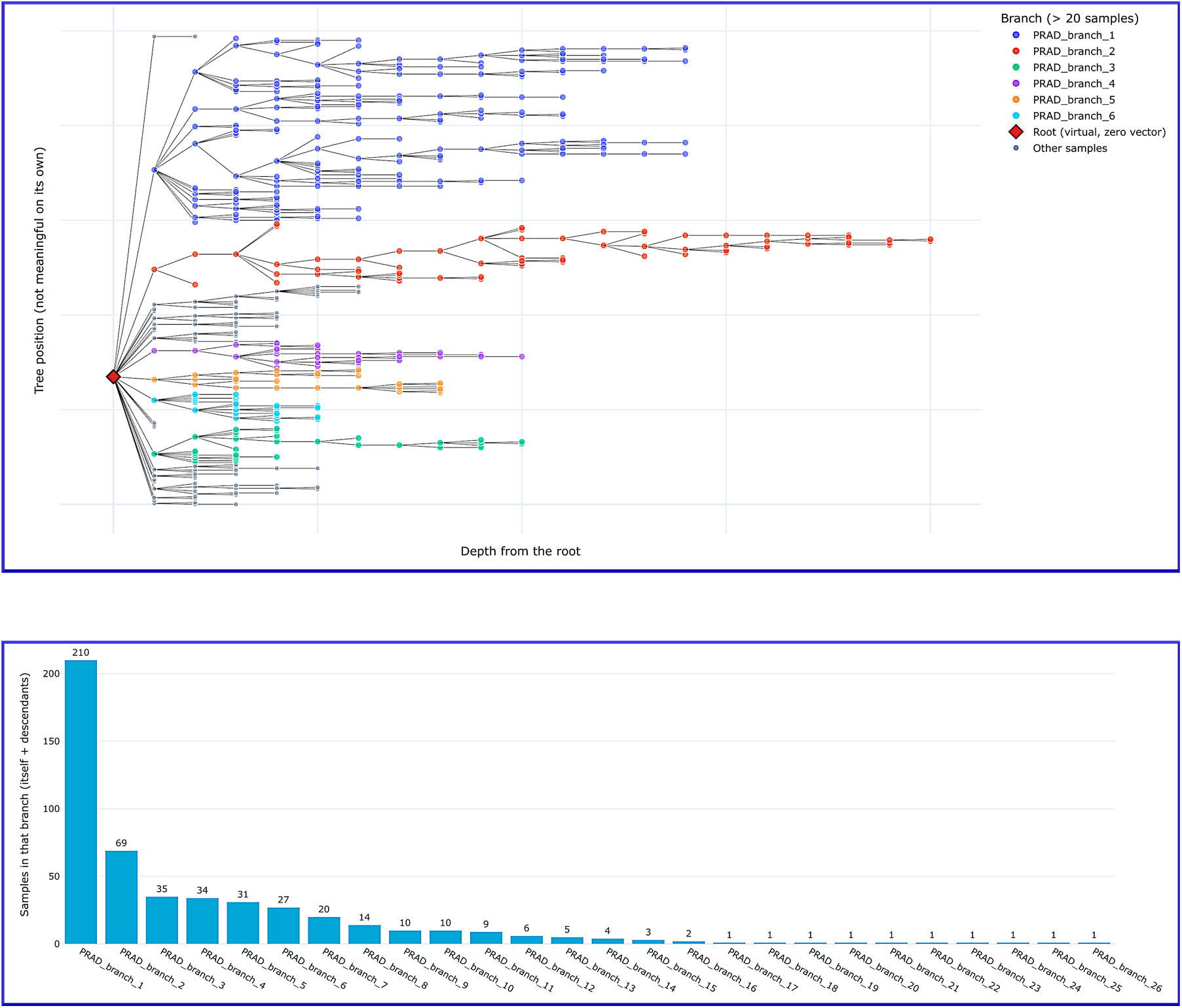
Taxonomy of PRAD tumors according to the MST algorithm. Upper panel: Branches in the MST scheme. The root, represented as a red point, is a fictitious sample with zero activated T-genes. The tree flows mainly in the direction of increasing active T-genes, showing that this number is a good temporal indicator. Lower panel: number of samples in each branch. There are 26 branches; the first six collect 406 of the 499 samples (81.4%), with sizes 210, 69, 35, 34, 31 and 27.

**Table 2.** Main pathway deregulation programs of the six molecular subtypes identified by the MST in PRAD tumors. For each subtype: size, dominant program, representative pathways, effective T frequency in the subtype, frequency in the global tumor population, and enrichment ratio (subtype/global). The six main branches contain 406 of the 499 PRAD tumors (81.4%); the remaining 93 samples are distributed across 20 minor branches (not characterized in detail due to limited statistical power).

| Subtype | Samples | Dominant program | Representative pathway | Freq. subtype | Freq. global | Enrichment |
|---|---|---|---|---|---|---|
| B1 | 210 (42.1%) | MET / NFE2L2 / energy metabolism | MET Receptor Activation | 0.66 | 0.56 | 1.18 |
|  |  |  | NFE2L2 regulating ER-stress genes | 0.61 | 0.55 | 1.1 |
|  |  |  | Utilization of Ketone Bodies | 0.58 | 0.5 | 1.16 |
| B2 | 69 (13.8%) | RAS / apoptosis / telomere | SMAC (DIABLO) binds to IAPs | 0.72 | 0.13 | 5.65 |
|  |  |  | Activation of caspases through apoptosome | 0.72 | 0.13 | 5.65 |
|  |  |  | Alternative Lengthening of Telomeres | 0.8 | 0.14 | 5.86 |
| B3 | 35 (7.0%) | PTEN / translation | Regulation of PTEN mRNA translation | 0.54 | 0.22 | 2.44 |
|  |  |  | ceRNAs regulate PTEN translation | 0.54 | 0.22 | 2.44 |
| B4 | 34 (6.8%) | DNA repair / lipid metabolism | Defective Mismatch Repair MSH6 | 0.71 | 0.18 | 3.87 |
|  |  |  | Glycerophospholipid catabolism | 0.74 | 0.36 | 2.07 |
| B5 | 31 (6.2%) | Inflammation / LXR lipid metabolism | The AIM2 inflammasome | 0.84 | 0.35 | 2.42 |
|  |  |  | NR1H2 & NR1H3 (LXR) lipolysis | 0.68 | 0.15 | 4.51 |
| B6 | 27 (5.4%) | Neuronal development / BER | Regulation of cortical dendrite branching | 0.67 | 0.43 | 1.56 |
|  |  |  | Defective Base Excision Repair NTHL1 | 0.52 | 0.31 | 1.68 |

#### Subtype B1 (210 samples, 42.1%): MET, NFE2L2, and energy metabolism

This is the largest subtype. Its most distinctive feature is the high frequency of MET receptor activation and its downstream effectors. MET activation is a known oncogenic driver in prostate cancer [25] and is associated with aggressive disease and resistance to androgen deprivation therapy. NFE2L2 (NRF2) deregulation controls the cellular response to oxidative stress and is increasingly recognized as a driver of metabolic reprogramming in cancer [26]. A shift in energy metabolism toward ketone body utilization and altered TCA cycle activity is consistent with the metabolic flexibility observed in advanced tumors [27]. The size of this subtype suggests that these functional features are shared by a large fraction of PRAD tumors, regardless of their genomic background.

#### Subtype B2 (69 samples, 13.8%): RAS, apoptosis, and telomere maintenance

This subtype is characterized by a striking deregulation of RAS signaling, apoptotic pathways, and telomere maintenance. Constitutive activation of RAS signaling is a hallmark of aggressive, treatment-resistant tumors. Although RAS mutations are rare in primary prostate cancer, the functional activation of this pathway has been associated with metastatic progression and resistance to therapy [28]. Deregulation of apoptosis, with a strong signature of both intrinsic and extrinsic apoptotic pathways, and activation of alternative telomere maintenance mechanisms are consistent with a mechanism of replicative immortality distinct from TERT promoter mutations.

#### Subtype B3 (35 samples, 7.0%): PTEN regulation and translation

This subtype is characterized by deregulation of PTEN translation and its regulation by competing endogenous RNAs, suggesting a post-transcriptional mechanism of PTEN loss rather than genomic deletion or mutation.

#### Subtype B4 (34 samples, 6.8%): DNA repair and lipid metabolism

This subtype is characterized by deregulation of DNA mismatch repair and lipid metabolism. Defective DNA mismatch repair is a known mechanism of genomic instability and is associated with a subset of prostate cancers, particularly those with SPOP mutations [29]. Altered lipid metabolism is increasingly recognized as a metabolic vulnerability in prostate cancer [30].

#### Subtype B5 (31 samples, 6.2%): Inflammation and LXR-mediated lipid metabolism

This subtype is characterized by inflammasome activation and LXR-mediated regulation of lipid metabolism. The AIM2 inflammasome is a key component of innate immunity and has been implicated in cancer progression through the release of pro-inflammatory cytokines [31]. Deregulation of LXR-mediated lipid metabolism controls cholesterol efflux, bile acid homeostasis, and triglyceride lipolysis, suggesting a metabolic reprogramming that favors lipid accumulation and may support tumor growth.

#### Subtype B6 (27 samples, 5.4%): Neuronal development and base excision repair

This subtype is characterized by deregulation of neuronal development pathways and base excision repair. Deregulation of neuronal development pathways, including MECP2, SLIT/ROBO, and cortical dendrite branching, is consistent with the known neuroendocrine differentiation that occurs in a subset of aggressive tumors [32,33,34].

### 3.4 Relationship between MST subtypes and known PRAD taxonomy

The functional subtypes identified by the MST may be compared with the genomic taxonomy established by the TCGA for PRAD [9]. The TCGA defined seven subtypes based on driver genomic events: fusions involving ERG (46%), ETV1 (8%), ETV4 (4%), and FLI1 (1%), mutations in SPOP (11%), FOXA1 (3%), and IDH1 (1%).

The six molecular subtypes identified by the MST reveal a previously unrecognized level of functional heterogeneity in PRAD.

The largest subtype (B1), characterized by MET activation, NFE2L2 deregulation, and altered energy metabolism, represents a functional program that is not captured by any single genomic driver. MET activation is known to occur in a subset of prostate cancers and is associated with aggressive disease, but it is not a defining feature of the TCGA subtypes. Similarly, NFE2L2 deregulation and metabolic reprogramming are common features of advanced tumors that cut across genomic categories. The size of this subtype (42% of tumors) suggests that these functional features are shared by a large fraction of PRAD tumors, regardless of their genomic background.

The second largest subtype (B2), characterized by RAS activation, apoptosis, and telomere maintenance, represents a functionally aggressive program that is not well captured by genomic classification. RAS mutations are rare in primary prostate cancer, but functional activation of the pathway, possibly through epigenetic or post-translational mechanisms, may drive a subset of tumors toward aggressive disease. The enrichment of telomere maintenance pathways in this subtype is particularly interesting, as it suggests a mechanism of replicative immortality that is distinct from the TERT promoter mutations commonly found in other cancers.

The four minor subtypes (B3–B6) each represent a distinct functional program that aligns with known biological features of prostate cancer. B3, with PTEN deregulation, aligns with the well-established role of PTEN loss in prostate cancer. B4, with DNA mismatch repair defects, aligns with the subset of tumors that have defective DNA repair and may benefit from immune checkpoint inhibitors. B5, with inflammasome activation and LXR-mediated lipid metabolism, aligns with the growing recognition of inflammation and metabolic reprogramming as drivers of prostate cancer. B6, with neuronal development and base excision repair, aligns with the neuroendocrine differentiation that occurs in a subset of aggressive tumors.

The correspondence between the MST subtypes and the TCGA genomic taxonomy is partial but informative. B3 and B4 align with specific genomic features (PTEN loss and mismatch repair defects). B5 may correspond to the triple-negative subtype. B2 does not have a genomic counterpart, suggesting that it represents a functional program that is not captured by genomic classification. B1, the largest, encompasses multiple genomic subtypes, suggesting that MET activation and metabolic reprogramming are common functional features that cut across genomic categories. B6 may correspond to neuroendocrine tumors, a rare but aggressive variant. This partial correspondence suggests that functional and genomic taxonomies are complementary rather than redundant.

### 3.5 Step 1 — Protection mechanisms in normal prostatic tissue

Figure 3, upper panel, shows the frequencies of active pathways in normal tissue. Notice that there are less than 1400 pathways, as compared with the more than 2400 pathways active in tumors. The global averages over the 52 normal prostatic samples identify the protection mechanisms maintained in the normal population. Table 3 summarizes these mechanisms, grouped by biological program, with representative pathways and their average effective N frequency.

**Figure 3.**
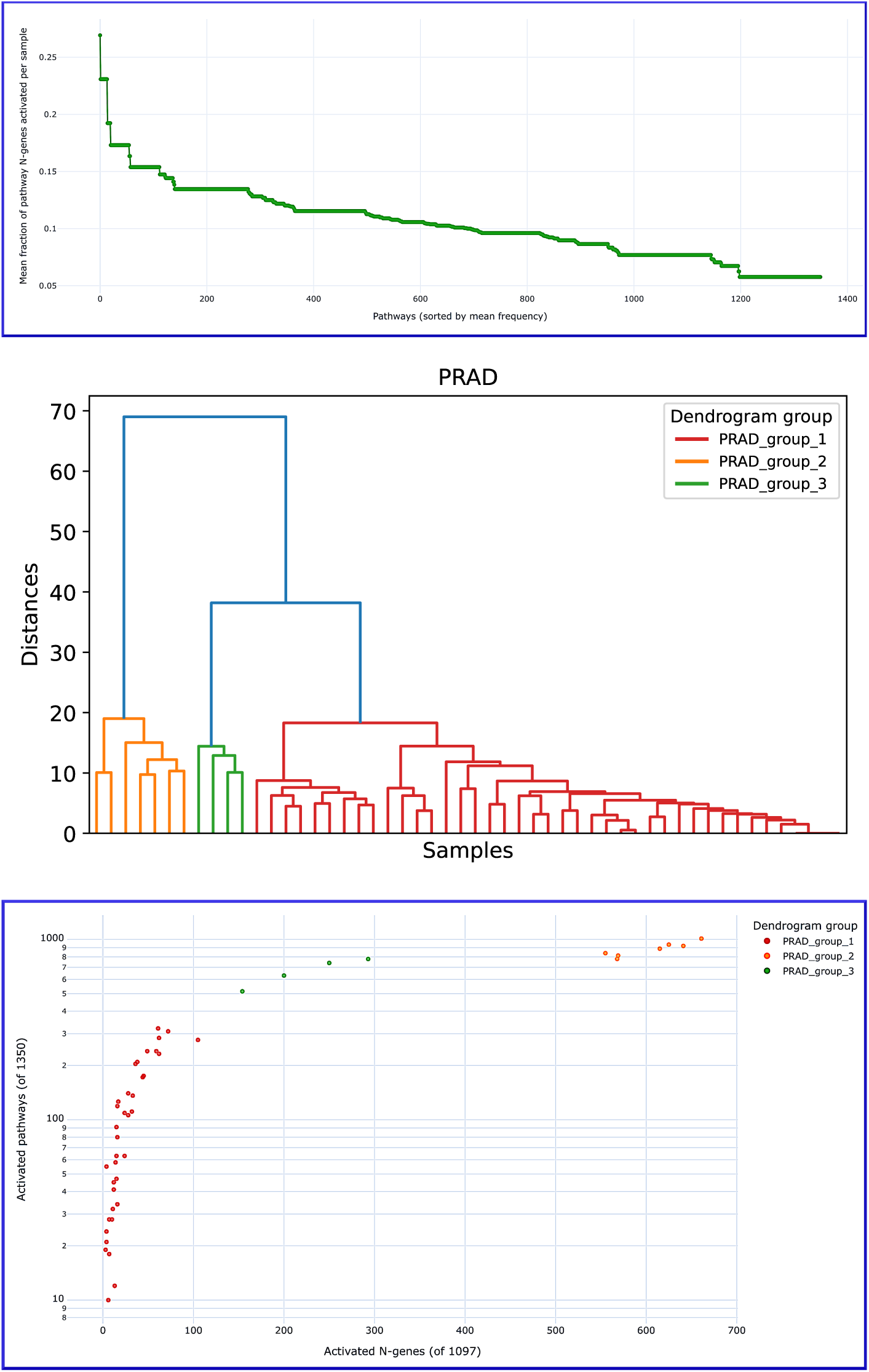
Clustering of normal samples according to the effective N frequency vectors in PRAD. Upper panel: mean frequency of pathways in normal prostatic samples, sorted by mean frequency (Step 1). Middle panel: dendrogram showing three groups of normal samples. Lower panel: number of active pathways as a function of the total number of active N-genes in each sample, with samples colored according to the clusters. The groups, ordered G2→G3→G1, represent stages of somatic evolution (Step 2).

**Table 3.** Protection mechanisms in normal prostatic tissue. For each biological program, representative pathways, average effective N frequency in the normal population, and inferred status (maintained, partially maintained, partially lost, or lost).

| **Protection program** | **Representative pathway** | **Avg. N freq.** | **Status** |
|---|---|---|---|
| Tie2 signaling | Tie2 Signaling | 0.27 | Maintained |
| FGFR2 signaling | FGFR2 ligand binding and activation | 0.23 | Maintained |
| Cell cycle regulation (APC/C) | APC/C-mediated degradation of cell cycle proteins | 0.17 | Partially maintained |
| DNA repair (BER) | Base Excision Repair | 0.13 | Partially maintained |
| TGF-beta signaling | Signaling by Activin | 0.21 | Partially maintained |
| FGFR1 signaling | FGFR1 ligand binding and activation | 0.15 | Partially maintained |
| IGF1R/Insulin signaling | IGF1R signaling cascade | 0.15 | Partially maintained |
| NFE2L2/KEAP1 | KEAP1-NFE2L2 pathway | 0.14 | Partially maintained |
| Integrin signaling | Integrin signaling | 0.14 | Partially maintained |
| ERBB4 signaling | Signaling by ERBB4 | 0.14 | Partially maintained |
| Apoptosis | Intrinsic Pathway for Apoptosis | 0.07 | Lost |
| Senescence | Cellular Senescence | 0.11 | Partially lost |
| Stress response | HSF1 activation | 0.08 | Lost |
| Homologous recombination repair | Homology Directed Repair | 0.11 | Partially lost |
| Telomere maintenance | Telomere Maintenance | 0.1 | Partially lost |
| Notch signaling | Signaling by NOTCH1 | 0.1 | Partially lost |
| WNT signaling | Signaling by WNT | 0.11 | Partially lost |
| RAS/MAPK signaling | RAF/MAP kinase cascade | 0.12 | Partially lost |
| Immune response | Interferon Signaling | 0.09 | Lost |

In order to analyze the Table, we take the highest observed frequencies (∼ 0.3) as an approximate ceiling of what a protection program can achieve in this cohort. The data reveal a clear hierarchy in the loss of protection. The mechanisms that are most robustly maintained in the normal population are the signaling pathways that control growth factor receptor activity (FGFR2, Tie2, FGFR1, IGF1R, ERBB4) and the cell cycle regulatory machinery (APC/C). These represent the first line of defense against uncontrolled proliferation. At an intermediate level, we find DNA repair (base excision repair), TGF-beta signaling, NFE2L2/KEAP1-mediated oxidative stress response, and integrin signaling. These mechanisms are partially maintained, suggesting that they are beginning to be lost in a fraction of the normal population. At the lowest level, we find the mechanisms that are most critical for preventing tumorigenesis: apoptosis, senescence, stress response, homologous recombination repair, telomere maintenance, Notch signaling, WNT signaling, RAS/MAPK signaling, and immune response. These are either lost or partially lost in the normal population, indicating that their inactivation is an early event in somatic evolution.

This hierarchy is consistent with the hallmarks of cancer [35,36]. The loss of apoptosis and senescence allows cells to survive and proliferate despite oncogenic stress. The loss of DNA repair and telomere maintenance leads to genomic instability. The loss of immune response allows the emerging tumor to evade immune surveillance. The maintenance of growth factor receptor signaling and cell cycle regulation until later stages may reflect the fact that these pathways are essential for normal tissue homeostasis and are only fully deregulated after the tumor is established.

### 3.6 Step 2 — Sequences of protection loss in normal prostatic tissue

Hierarchical clustering of the 52 normal prostatic samples on their effective N frequency vectors identifies three groups, as shown in the dendrogram of Figure 3, central panel. The groups may be ordered by the number of active N-genes, from least to most advanced somatic evolution, as **G2 → G3 → G1** (Figure 3, lower panel): G2 contains the samples with the largest number of active N-genes and therefore the most intact protection, G3 is intermediate, and G1 contains the samples with the fewest active N-genes and the most advanced loss of protection. The groups are highly unbalanced: G1 contains 41 of the 52 samples, whereas G2 contains 7 and G3 contains 4.

To ensure consistency with the global averages of Step 1 (Table 3), we use the same criterion to classify protection in each group: a group average of **≥ 0.3** (the approximate ceiling observed among the most robustly maintained programs in the cohort, such as Tie2 signaling) indicates maintained protection, whereas a group average **< 0.3** indicates loss of protection. The **onset of loss** is defined as the first group, in the order G2 → G3 → G1, in which the pathway falls below 0.3. Table 4 reports, for all the protection programs of Table 3, the group averages and the onset of loss.

**Table 4.** Loss of protection along the ordered normal groups in PRAD. Protection is considered maintained when the group average is ≥ 0.3 (the approximate ceiling observed in the cohort) and lost when it falls below 0.3. The onset of loss is the first group, in the order G2 → G3 → G1, in which the pathway falls below 0.3. This table includes all the protection programs listed in Table 3.

| Protection program | Representative pathway | G2 | G3 | G1 | Onset of loss | Trajectory |
|---|---|---|---|---|---|---|
| **Early loss (G2)** |  |  |  |  |  |  |
| Stress response | HSF1 activation | 0.14 | 0 | 0.07 | G2 | Loss |
| RTK (FGFR2) | FGFR2 ligand binding and activation | 0 | 0.75 | 0.23 | G2 | Transient peak (G3) |
| Integrin signaling | Integrin signaling | 0 | 1 | 0.09 | G2 | Transient peak (G3) |
| Notch signaling | Signaling by NOTCH1 | 0.23 | 0.55 | 0.04 | G2 | Transient peak (G3) |
| RAS/MAPK signaling | RAF/MAP kinase cascade | 0.28 | 0.53 | 0.05 | G2 | Transient peak (G3) |
| **Intermediate loss (G3)** |  |  |  |  |  |  |
| Cell cycle regulation (APC/C) | APC/C-mediated degradation of cell cycle proteins | 0.71 | 0 | 0.1 | G3 | Loss |
| NFE2L2/KEAP1 | KEAP1-NFE2L2 pathway | 1 | 0.13 | 0 | G3 | Loss |
| DNA repair (BER) | Base Excision Repair | 0.81 | 0.25 | 0 | G3 | Loss |
| Homologous recombination repair | Homology Directed Repair | 0.71 | 0.1 | 0 | G3 | Loss |
| Apoptosis | Intrinsic Pathway for Apoptosis | 0.5 | 0 | 0 | G3 | Loss |
| Telomere maintenance | Telomere Maintenance | 0.67 | 0.17 | 0 | G3 | Loss |
| TGF-beta signaling | Signaling by Activin | 1 | 0.25 | 0 | G3 | Loss |
| Senescence | Cellular Senescence | 0.63 | 0.18 | 0 | G3 | Loss |
| Immune response | Interferon Signaling | 0.35 | 0.27 | 0.03 | G3 | Loss |
| **Late loss (G1)** |  |  |  |  |  |  |
| RTK (Tie2) | Tie2 Signaling | 1 | 1 | 0.22 | G1 | Loss |
| RTK (FGFR1) | FGFR1 ligand binding and activation | 0.43 | 0.56 | 0.05 | G1 | Loss |
| IGF1R/Insulin signaling | IGF1R signaling cascade | 0.33 | 0.58 | 0.07 | G1 | Loss |
| ERBB4 signaling | Signaling by ERBB4 | 0.5 | 0.5 | 0.05 | G1 | Loss |
| WNT signaling | Signaling by WNT | 0.4 | 0.43 | 0.03 | G1 | Loss |

*Note:* Pathways marked "Transient peak (G3)" have a group average ≥ 0.3 only in G3, which contains 4 samples. These are reported as exploratory observations rather than stable trajectory classes, because they may reflect cell-type composition in G3 rather than a reproducible loss–recovery pattern. The three classes shown exhaust the patterns present in the data.

Three trajectories are distinguishable (Table 3b). First, a group of programs shows an **early onset of loss in G2**, with a group average below 0.3 already in the least advanced group: the stress response (HSF1 activation) is the clearest case, and four additional programs—FGFR2, integrin, Notch, and RAS/MAPK signaling—are also below 0.3 in G2, but rise above 0.3 in G3 and fall below it again in G1, a transient pattern that is reported as exploratory (see note). Second, a group of programs shows an **intermediate onset of loss in G3**: their group average is at or above 0.3 in G2, falls below 0.3 in G3, and remains below it in G1. This group includes the cell cycle regulatory machinery (APC/C), the NFE2L2/KEAP1 oxidative-stress pathway, base excision repair, homologous recombination repair, the intrinsic pathway of apoptosis, telomere maintenance, TGF-beta signaling, cellular senescence, and interferon signaling. Third, a group of programs shows a **late onset of loss in G1**: their group average remains at or above 0.3 through G3 and falls below 0.3 only in G1. This group includes Tie2 signaling, FGFR1, IGF1R, ERBB4, and WNT signaling.

The hierarchy of Table 3 reflects the final state of the cohort (dominated by G1), whereas the group-level sequence of Step 2 reveals that most of the loss occurs earlier, in G3. Programs that Table 3 classifies as “partially maintained” (e.g., TGF-beta signaling, base excision repair) are in fact lost by G3 when compared to the 0.3 ceiling, and programs that Table 3 classifies as “lost” (e.g., apoptosis, senescence) are lost even earlier than their final frequencies suggest. The group-level sequence therefore complements the global averages of Step 1.

A caveat follows. The groups are highly unbalanced: G1 contains 41 of the 52 samples, whereas G2 contains 7 and G3 contains 4. The G2 and G3 averages rest on very few samples, so a value of 0.00 or 1.00 in these groups may reflect sampling noise rather than a genuine biological state. This is particularly relevant for the transient class, whose G3 values rest on 4 samples. The group-level comparison should therefore be read as a qualitative, complementary view rather than as an independent confirmation of Step 1. We report the group-level sequence with this caveat, and we do not use it to reorder the protection hierarchy of Table 3.

### 3.7 Step 3 — Taxonomy of normal prostatic tissue

The MST was also applied to the 52 normal prostatic samples, using the same distance metric and the same pathway frequency vectors as in Step 3 for tumors. In contrast to the tumor compartment, the MST built on normal samples does not subdivide the normal population into distinct branches: all 52 samples remain connected in a single branch. This result is consistent with the view that the normal prostatic compartment, as represented in TCGA, is transcriptionally more homogeneous than the tumor compartment, at least at the level of pathway deregulation captured by the N/T framework. Alternatively, the absence of subdivision may reflect the limited size of the normal cohort (n=52) rather than a biological absence of substructure, and we cannot separate these two contributions with the current data. Because the MST does not subdivide the normal population, the normal compartment is characterized through the global averages of Step 1 (Table 3) and the group-level sequence of Step 2 (Section 3.6).

### 3.8 Comparison between protection loss in normals and progression gain in tumors

The comparison between the protection mechanisms lost in normals (Table 3) and the progression programs gained in tumors (Table 2) reveals a striking complementarity. Subtype B4, characterized by DNA mismatch repair deregulation in tumors, corresponds to the loss of DNA repair protection in normals. Subtype B3, characterized by PTEN deregulation in tumors, corresponds to the partial maintenance of IGF1R/Insulin signaling protection in normals. Subtype B2, characterized by RAS activation, apoptosis, and telomere maintenance in tumors, corresponds to the loss of apoptosis protection and telomere maintenance protection in normals. Subtype B1, characterized by MET activation and NFE2L2 deregulation in tumors, corresponds to the partial maintenance of NFE2L2/KEAP1 protection in normals. Subtype B5, characterized by inflammation and LXR-mediated lipid metabolism in tumors, corresponds to the loss of immune response protection in normals. Subtype B6, characterized by neuronal development and NTHL1 base excision repair in tumors, corresponds to the partial maintenance of base excision repair protection in normals.

This complementarity strongly supports the N/T framework: the loss of N-gene activity in normal tissue (somatic evolution) precedes and predisposes to the gain of T-gene activity in tumors (tumor progression). The two phases of carcinogenesis are not independent but are linked by the specific mechanisms that are lost and gained.

### 3.9 Verification with complete gene panels

#### 3.9.1 T-gene panel and MST subtypes

The eight T-genes of the perfect panel were computed per MST branch (Table 4). The panel genes reproduce the six subtypes, and identify the gene or genes that most clearly characterize each one.

**Table 4.** Activation of the perfect T-gene panel across MST branches in PRAD. For each gene, average activation per branch; the maximum is marked in bold.

| **Gene** | **B1** | **B2** | **B3** | **B4** | **B5** | **B6** | **Max** |
|---|---|---|---|---|---|---|---|
| EPHA10 | 0.79 | 0.65 | 0.63 | **0.85** | **0.87** | **0.85** | B4/B5/B6 |
| UCN | 0.66 | **0.97** | 0.66 | 0.32 | 0.32 | 0.63 | B2 |
| TMEM86A | **0.6** | 0.41 | 0.34 | 0.35 | **0.58** | 0.44 | B1/B5 |
| RPL26P23 | 0.5 | 0.28 | **0.54** | **0.59** | **0.55** | 0.52 | B3/B4/B5 |
| RBM15B | 0.47 | 0.15 | **0.6** | 0.5 | 0.52 | 0.52 | B3 |
| NPY4R2 | 0.28 | 0.01 | 0.26 | 0.35 | **0.58** | 0.19 | B5 |
| TEKT4 | 0.22 | **0.45** | 0.2 | 0.27 | 0.13 | 0.26 | B2 |
| ENSG00000271895 | 0.2 | 0.35 | **0.43** | 0.06 | 0.03 | 0.22 | B3 |

TMEM86A marks B1; UCN and TEKT4 mark B2 (RAS/apoptosis/telomere); the lncRNA ENSG00000271895, RPL26P23 and RBM15B mark B3 (PTEN); EPHA10 and RPL26P23 marks B4 (DNA repair); EPHA10, TMEM86A, RPL26P23 and NPY4R2 mark B5 (inflammation/LXR); EPHA10 marks B6 (Neuronal development/base excision repair). This confirms the coherence between the pathway-level view (Table 2) and the gene-level view.

The trajectory of these genes along the clustering groups (G2→G1→G3→G4) is heterogeneous: UCN increases with progression (onset G3), TEKT4 shows a late gain (G4), EPHA10 and TMEM86A peak early (G1), RBM15B peaks in G3, and RPL26P23 and NPY4R2 decline in G4.

This heterogeneity is consistent with the idea that the panel captures subtype-specific programs rather than a single monotonic progression axis.

#### 3.9.2 N-gene panel and the hierarchy of protection

The activation of the eleven N-genes of the perfect panel was computed in the normal PRAD population. Because the MST yields a single normal group, the average is over the whole normal population (Table 5).

**Table 5.** Activation of the perfect N-gene panel in normal PRAD tissue. For each gene, average N frequency and associated protection mechanism (from Table 3).

| **Gene Symbol** | **N frequency** | **Associated protection mechanism** | **Status in Table 3** |
|---|---|---|---|
| SEPTIN10 | 0.31 | Cell cycle regulation (APC/C) / chromatid cohesion | Partially maintained (0.173) |
| CA14 | 0.19 | Ion homeostasis / transport | Not listed explicitly |
| TES | 0.15 | Cell–cell adhesion / integrin signaling | Partially maintained (0.144) |
| PARM1 | 0.15 | Differentiation / luminal identity | Not listed explicitly |
| APOBEC3C | 0.14 | Immune response / DNA repair (BER) | Lost (0.094) / partially maintained (0.128) |
| ENSG00000224984 | 0.14 | Unknown (lncRNA) | – |
| TMT1A | 0.12 | Fatty acid metabolism / lipid metabolism | Not prominent in PRAD |
| SLC18A2 | 0.1 | Neurotransmitter transport / neuronal protection | Not prominent in PRAD |
| SNORD104 | 0.08 | rRNA processing / translation | Not listed |
| RNA5SP342 | 0.08 | rRNA processing / translation | Not listed |
| HSPA1L | 0.06 | Stress response (HSF1 activation) | Lost (0.077) |

The normal panel genes reproduce the hierarchical loss of protection described in Section 3.5:

SEPTIN10 (0.308), a regulator of cytokinesis and the cytoskeleton, has the highest N frequency, consistent with the robust maintenance of cell cycle regulation (APC/C, 0.173) as the first line of defense against uncontrolled proliferation. CA14 (0.192) and TES (0.154) follow. CA14 is involved in ion homeostasis, and TES is a tumor suppressor linked to cell adhesion and integrin signaling (0.144). Their relatively high frequencies indicate that protection of epithelial architecture and ion balance is maintained in normal tissue. PARM1 (0.154) and APOBEC3C (0.135) reflect the protection of luminal differentiation and immune/mRNA editing mechanisms, respectively. APOBEC3C links the immune response (0.094) and DNA repair (BER, 0.128) protection programs. TMT1A (0.115), SLC18A2 (0.096), SNORD104 (0.077), RNA5SP342 (0.077), and HSPA1L (0.058) have the lowest frequencies, consistent with the early loss of lipid metabolism, neurotransmitter transport, rRNA processing, and stress response protection in the normal population. HSPA1L, a chaperone of the HSP70 family, is the gene with the lowest N frequency, matching the early loss of HSF1-mediated stress response protection (0.077) described in Table 3.

Summarizing, the complete gene panels reproduce the main subtypes and the hierarchy of protection mechanisms identified by the N/T framework at the pathway level. The T-genes capture the functional programs of the six MST subtypes, including several unanticipated possible associations (e.g., neuroendocrine signaling in the RAS subtype, nucleolar stress in the DNA repair subtype) that enrich the taxonomy. The N-genes capture the hierarchical loss of protection, with cell cycle and adhesion genes showing the highest frequencies and stress response and rRNA processing genes showing the lowest. This consistency between the pathway-level and gene-level views strengthens the validity of the N/T framework and supports the interpretation of the MST subtypes and protection profiles as biologically meaningful entities.

### 3.10. Cross-cancer comparison

The analysis of liver hepatocellular carcinoma (LIHC), lung adenocarcinoma (LUAD), and glioblastoma (GBM) follows the same three-step logic applied to PRAD. In this section we draw the global picture across the four tissues; the extended per-tissue analyses are provided in the Supplementary Material. Table 6 summarizes the molecular taxonomy, and Table 7 summarizes the protection mechanisms maintained in normal tissue.

#### 3.10.1 Molecular taxonomy

**Table 6.** Molecular taxonomy across PRAD, LIHC, LUAD, and GBM tumors.

| **Feature** | **PRAD** | **LIHC** | **LUAD** | **GBM** |
|---|---|---|---|---|
| Main tumor subtypes (MST) | 6 | 2 | 1 dominant group | 1 dominant group |
| Dominant subtype frequency | 0.42 (210 samples) | 0.85 (320 samples) | 1 (533/535 samples) | 1 (169/169 samples) |
| Main tumor programs | MET/NFE2L2, RAS/apoptosis/telomere, PTEN, DNA repair, inflammation/LXR, neuronal/NTHL1 | NTRK, senescence/LXR, phagosome/solute transport, selenium | Surfactant, FGFR1–4, glycosylation, selenium, cell cycle | TP53 loss, senescence evasion, cell cycle (G2/M), lipid metabolism, N-glycosylation, RTKs (NTRK2, FLT3, PDGFR), DNA repair deregulation |
| Correspondence with genomic taxonomy | Partial (PTEN, MMR) | Partial (proliferative/non-proliferative) | Partial (three transcriptional subtypes) | Partial (classical/proneural/mesenchymal; TP53, RTK axes) |

#### 3.10.2 Protection mechanisms in normal tissues

**Table 7.** Protection mechanisms across PRAD, LIHC, LUAD, and GBM.

| Mechanism | PRAD | LIHC | LUAD | GBM |
|---|---|---|---|---|
| DNA repair | Partially maintained (BER 0.13) | Maintained (BER 0.78) | Partially maintained (BRCA1 0.66, NEIL3 0.75) | Maintained (HR 0.80; BER 0.30) |
| Apoptosis | Lost (0.07) | Partially maintained (0.22–0.57) | Partially maintained (0.30–0.52) | Maintained (BAX 0.80) |
| Senescence | Lost (0.11) | Partially maintained (0.32–0.39) | Partially maintained (0.44–0.56) | Maintained (p16INK4A 1.00) |
| Immune response | Lost (0.09) | Partially maintained (0.20–0.36) | Partially maintained (0.35–0.65) | Partially maintained (phagosome 0.80) |
| Cell cycle | Partially maintained (0.17) | Partially maintained (0.50–0.57) | Maintained (0.47) | Maintained (CDK4/6 1.00) |
| RTKs | Maintained (Tie2 0.27) | Maintained (RTKs 1.0 in branch B2) | Maintained (FGFR2 0.57, MAPK 0.71) | Maintained (FGFR1 1.00, NTRK2 0.80) |

#### 3.10.3 Shared and tissue-specific features

**Shared across all four tissues:**

- DNA repair (BER, HR). DNA repair is present as a protection program in normal tissue in all four cohorts, although its maintenance status along progression differs: it is partially maintained in PRAD (Table 3) and maintained at higher frequency in LIHC, LUAD, and GBM (Table 7). The specific repair pathways involved differ across tissues. In tumors, two situations must be distinguished: in some tumors DNA repair does not appear among the dominant programs, which corresponds to the loss of the normal protective function; in others, DNA repair pathways appear with T-genes active, which reflects an aberrant transcriptional program.
- Cell cycle regulation is maintained in normal tissue, though to different degrees (lowest in post-mitotic brain).
- RTK signaling is protected in normals, though the specific RTKs differ (Tie2/FGFR in PRAD, RTK network in LIHC, FGFR2/MAPK in LUAD, FGFR1/NTRK2 in GBM). In tumors, RTK pathways become deregulated (T-genes active).
- The hierarchical loss of protection is observed where data are available: apoptosis and senescence are lost early (except in GBM normals, where they are maintained), DNA repair at an intermediate stage, and cell cycle and RTKs later.

**Tissue-specific:**

- **PRAD:** MET, NFE2L2, ketone bodies, Tie2.
- **LIHC:** NTRK, LXR/gluconeogenesis, phagosome acidification, solute transport.
- **LUAD:** surfactant, autophagy, hypoxia (HIF), NOTCH, extensive glycosylation, MAPK/RAS/RAF, FGFR2.
- **GBM:** TP53 loss (tumor), senescence evasion, intense lipid metabolism (cardiolipin, sphingolipids), N-glycosylation defects (ALG3), RTKs (NTRK2, FLT3, PDGFR), DNA repair deregulation, and high protection of apoptosis/senescence in normals.

#### 3.10.4 Connection between protection loss and tumor progression

In all four tissues, the loss of protection mechanisms in normals is reflected in the gain of corresponding progression programs in tumors. In PRAD, loss of apoptosis and telomere protection corresponds to the RAS/apoptosis/telomere tumor subtype. In LIHC, loss of senescence and NTRK protection corresponds to senescence evasion and NTRK activation in the dominant tumor subtype. In LUAD, loss of surfactant, MAPK, and FGFR2 protection corresponds to the deregulation of these programs in tumors. In GBM, the maintenance of apoptosis (0.80) and senescence (1.00) protection in normals contrasts with the near-universal TP53 loss (0.97) and senescence evasion (0.40) in tumors, indicating that loss of these protections is a key event. The maintenance of RTK protection in normals (FGFR1 1.00, NTRK2 0.80) contrasts with high RTK deregulation in tumors (NTRK2 0.90), suggesting that loss of RTK protection precedes RTK-driven progression. The maintenance of HR protection in normals (0.80) contrasts with the strong deregulation of DNA repair pathways in tumors (BRCA2 0.99, NEIL3 0.99), indicating that the normal repair protection is lost and replaced by an aberrant transcriptional program. The partial loss of hypoxia protection in normals (0.51) may predispose to the hypoxic adaptation observed in GBM. This supports the central claim of the N/T framework: the loss of N-gene activity in normal tissue (somatic evolution) precedes and predisposes to the gain of T-gene activity in tumors (tumor progression).

#### 3.10.5 Heterogeneity and tumor evolution

The number of main tumor subtypes identified by the MST decreases progressively from PRAD (6 subtypes) to LIHC (2 subtypes) to LUAD (1 dominant group) to GBM (1 dominant group). This gradation is paralleled by a decrease in the number of genes in the complete panel (from PRAD to GBM) [2], and by an increase in the number of activated T-genes (from PRAD to GBM) [4]. These observations may suggest that the molecular heterogeneity of the tumor decreases as the tumor becomes more aggressive and evolutionary advanced. In less aggressive tumors (PRAD), multiple alternative trajectories coexist, reflecting different combinations of driver events and pathway deregulations. In more aggressive tumors (LUAD, GBM), the tumor converges toward a common progression program that includes cell cycle deregulation, TP53 loss, metabolic reprogramming, and DNA repair deregulation, leaving little room for alternative trajectories. This convergence is not a failure of the method, but a biological property of advanced tumors: the selective pressures of the tumor microenvironment and the accumulation of driver events reduce the diversity of viable molecular states. The N/T framework captures this gradation naturally.

## 4. Discussion

### 4.1 Advantages of the proposed method

The proposed method has three advantages over existing pathway-based approaches. First, it eliminates dilution by non-informative genes: only N- and T-genes contribute to the signal, so a pathway is scored by the genes that actually drive the normal-to-tumor transition rather than by all annotated genes. Second, it provides a direct, normalized, and sample-by-sample measure of pathway deregulation—the effective frequency—which is intuitive, comparable across pathways and cohorts, and suitable for classification and clustering. Third, it allows separate evaluation of loss of protection in normal samples and gain of progression in tumor samples, providing a dual perspective on carcinogenesis that is not available in methods that analyze tumors alone.

The explicit three-step analytical logic—global averages, clustering-based sequences, and MST-based taxonomy—is a further methodological contribution. By separating the three levels of aggregation, the method avoids a common pitfall: conflating classification with progression. The MST is built on pathway frequency vectors, not on the number of activated genes, so the resulting branches are a genuine taxonomy; the fact that they correlate with the number of activated genes is a validation, not a circularity.

Beyond this, the most original contribution of this work is the joint analysis of normal and tumor compartments across four cancer types. Previous pathway-based taxonomies were developed and validated within a single tumor type. By applying the same three-step logic to PRAD, LIHC, LUAD, and GBM, and by analyzing both compartments in each tissue, we show that the loss of protection in normal tissue and the gain of progression programs in tumors share a common architecture across tissues, while the specific pathways involved are tissue-specific. This cross-cancer dimension is what allows the N/T decomposition of carcinogenesis to be tested as a general principle rather than as a description of a single cohort.

### 4.2 Limitations

The method has several limitations. It depends on pathway annotation: only about eleven thousand genes are annotated in Reactome, so a fraction of the N- and T-genes identified in each cohort cannot be mapped to pathways. This may bias the analysis toward well-studied pathways, which tend to have more annotated genes and therefore more opportunities to contain N- or T-genes. The method is also dataset-dependent, as N- and T-genes are specific to each cancer type and cohort, and it relies on significance thresholds that must be re-derived for each cohort. The dynamic interpretation depends on the ergodic assumption—that the ordered groups along the number of active genes can be read as a temporal sequence—which is an approximation.

A specific limitation concerns the group-level sequences in normal tissue. As reported in Section 3.6, the trajectories of several N-genes and protection pathways across G2 → G3 → G1 are not monotonic, showing transient peaks. This heterogeneity means that the hierarchy of protection is supported more strongly by the global averages of Step 1 (Table 3) than by the group-level sequence of Step 2. The three groups of normal samples may reflect biological substructure rather than a clean temporal sequence, and future work with larger normal cohorts should address this.

A further limitation concerns the cross-cancer comparison. The four cohorts differ widely in size and in the availability of normal samples—from 499 tumors and 52 normals in PRAD to very few normals in GBM—so differences in the number of subtypes identified across tissues may reflect differences in statistical power as much as differences in underlying biological heterogeneity. The pattern reported in Section 3.10.5 (six subtypes in PRAD, two in LIHC, one dominant program in LUAD and GBM) is therefore presented as consistent with, but not proof of, a contraction of molecular heterogeneity in more advanced tumors. Separating these two contributions would require matched cohort sizes or resampling analyses.

Finally, the number of activated genes is an indicator of temporal evolution, not a clinical staging tool. We emphasize that it should not be used for clinical decision-making without further validation.

### 4.3 Biological implications

The method has several potential biological implications. It could identify protection pathways that become deregulated early in somatic evolution, providing targets for early intervention, and progression pathways that become activated in advanced tumors, providing targets for therapy. In PRAD, the six molecular subtypes suggest testable hypotheses: the MET/NFE2L2 subtype may be sensitive to MET or NRF2 inhibitors, the RAS/apoptosis subtype to RAS-MAPK or apoptosis-inducing agents, the PTEN subtype to PI3K/AKT inhibitors, the DNA repair subtype to PARP inhibitors or immune checkpoint inhibitors, the inflammation/LXR subtype to anti-inflammatory agents or LXR agonists, and the neuronal development subtype to therapies targeting neuroendocrine differentiation. These hypotheses are speculative and require clinical validation, but they illustrate the potential of the method for guiding therapeutic decisions.

The hierarchical loss of protection in normal tissue also has implications for cancer prevention. The early loss of apoptosis, senescence, and immune response suggests that interventions that restore these mechanisms could prevent the emergence of tumors. The intermediate loss of DNA repair suggests that interventions that enhance DNA repair capacity could delay the accumulation of mutations. The late loss of growth factor receptor signaling and cell cycle regulation suggests that these pathways are less suitable for prevention but may be effective targets for therapy in advanced tumors.

### 4.4 Future work

Future work should proceed along four lines. First, the molecular subtypes identified here should be validated in independent cohorts and, where available, against clinical outcomes. Second, the method should be extended to single-cell RNA-seq data to resolve intratumoral heterogeneity, and integrated with gene deregulation networks such as those described by Castel et al. [24]. Third, the pathway analysis should be compared across Reactome and other databases such as KEGG, WikiPathways, and Gene Ontology, to assess the robustness of the subtypes to annotation choice. Fourth, the framework should be applied to additional cancer types and to the quantification of drug response in cell lines, and the use of the number of activated genes as a progression indicator should be explored in clinical research settings.

## 5. Conclusions

We propose a reformulation of pathway analysis based on effective activation frequencies of N- and T-genes, organized in three explicit steps—global averages, clustering-based sequences, and MST-based taxonomy—and apply it to construct a molecular taxonomy and progression description of tumors across four cancer types, using the full set of Reactome pathways. The method eliminates dilution by non-informative genes, allows direct quantification of pathway deregulation, facilitates sample classification, and unifies taxonomy with a molecular indicator of progression.

In PRAD, taken as the detailed case study, the MST resolves six molecular subtypes with distinct pathway deregulation profiles, which together account for 81.4% of the 499 tumors (Section 3.3). Analysis of the normal prostatic compartment reveals a hierarchy of protection loss: immune response, apoptosis, and senescence are lost or partially lost early, DNA repair at an intermediate stage, and cell cycle regulation and RTK signaling last. The perfect T and N gene panels reproduce the subtype structure and the protection hierarchy at gene level, connecting the pathway-level and gene-level views.

The most distinctive contribution of this work emerges from the cross-cancer comparison. DNA repair, cell cycle, and RTK protection are maintained in normal tissue across PRAD, LUAD, LIHC, and GBM, whereas the specific pathways lost in normals and gained in tumors are tissue-specific (MET/NFE2L2 in PRAD; NTRK/LXR/phagosome in LIHC; surfactant/FGFR/glycosylation in LUAD; TP53 loss/lipid metabolism/glycosylation in GBM). This shared architecture—loss of protection in normal tissue preceding and predisposing to gain of progression programs in tumors— supports the N/T decomposition of carcinogenesis into somatic evolution and tumor progression as a general principle rather than a tissue-specific observation. The number of molecular subtypes identified by the MST decreases from PRAD (six) to LIHC (two) to LUAD and GBM (one dominant program), a pattern consistent with a contraction of molecular heterogeneity as tumors become more evolutionary advanced, although the present data do not allow this trend to be separated from differences in cohort size and sampling.

The method is general and applicable across cancer types, although quantitative results depend on the underlying cohort, on Reactome annotation coverage, and on thresholds that must be re-derived for each dataset. The number of activated genes is an indicator of temporal evolution, not a clinical staging tool, and should not be used for clinical decision-making without further validation. Future work should extend the framework to single-cell data, to other pathway databases, and to the quantification of drug response in cell lines, and should test the clinical relevance of the molecular subtypes identified here.

## Data and code availability statement

All data used in this study are publicly available through the TCGA Research Network (https://www.cancer.gov/tcga) and the Reactome website (www.https://reactome.org/). Gene classification and pathway processing were performed using the in-house code available at https://github.com/JoanANievesCuadrado/pathways-ntgenes.

## Ethical Considerations

TCGA data are public, so no ethical approval is required. Reactome and TCGA licenses are respected.

