## Supplementary Material for "N- and T-genes framework for pathway deregulation: Taxonomy, progression, and protection in cancer"

#### Cross-cancer comparison of the N/T framework

This Supplementary Material presents the detailed results for liver hepatocellular carcinoma (LIHC), lung adenocarcinoma (LUAD), and glioblastoma (GBM), complementing the main text analysis of prostate adenocarcinoma (PRAD). For each cancer type, we describe the progression mechanisms of tumors, the protection mechanisms in normal tissue, and the connection between the two. A cross-cancer comparison that highlights shared and tissue-specific features is presented in the main text..

**Note on the three-step logic.** As in the main text, each tissue is analyzed in three explicit steps: (i) global cohort averages (Level 1); (ii) averages over the groups of hierarchical clustering, ordered by the number of active genes (Level 2); (iii) MST taxonomy and branch averages (Level 3). For GBM, the clustering groups overlap and the MST collapses into a single branch; therefore, GBM is analyzed **only at Level 1 (global)**, and this exception is documented explicitly in section S3.

---

#### S1. Liver Hepatocellular Carcinoma (LIHC)

##### S1.1 Level 1 — Pathways important in LIHC tumors (global)

**Supplementary Table 1.** Global averages over the LIHC tumor cohort identify the dominant programs. Representative pathways, grouped by mechanism:

| Mechanism | Representative pathway | Global freq. |
|---|---|---|
| Neurotrophic signaling | NTF3 activates NTRK2 (TRKB) signaling | 0.88 |
| Neurotrophic signaling | NTF3 activates NTRK3 signaling | 0.88 |
| Senescence evasion | Evasion of Oncogene Induced Senescence Due to p14ARF Defects | 0.8 |
| Senescence evasion | Evasion of Oxidative Stress Induced Senescence Due to p14ARF Defects | 0.8 |
| DNA synthesis | Plus-strand DNA synthesis | 0.8 |
| DNA synthesis | Minus-strand DNA synthesis | 0.8 |
| HIV / reverse transcription | Reverse Transcription of HIV RNA | 0.8 |
| Glycosylation | Defective MOGS causes CDG-2b | 0.73 |
| Glycosylation | Defective ALG3 causes CDG-1d | 0.72 |
| Cell cycle (G2/M) | Phosphorylation of proteins involved in the G2/M transition by Cyclin A:Cdc2 complexes | 0.73 |
| Immune evasion | Blockage of phagosome acidification | 0.74 |
| DNA repair (MMR) | Defective Mismatch Repair Associated With MSH6 | 0.68 |
| Lipid metabolism (LXR) | NR1H2 & NR1H3 regulate gene expression linked to gluconeogenesis | 0.66 |
| Selenium | Methylation of MeSeH for excretion | 0.73 |

| | | |
|---|---|---|
| metabolism | | |
| Solute transport | Defective SLCO2B3 causes hyperbilirubinemia, Rotor type (HBLRR) | 0.65 |

These global values constitute the reference for the sequence (S1.2) and taxonomy (S1.3) analyses.

#### S1.2 Level 2 — Sequences of pathways activation in LIHC tumors

Hierarchical clustering of LIHC tumor samples on their effective T frequency vectors identifies **four groups**, ordered by the number of active T-genes as **G4** → **G2** → **G1** → **G3**. Comparing pathway averages across these ordered groups reveals a staggered gain of programs.

**Criterion:** active if group frequency > global; appreciable change if |Δ| ≥ 0.10 between consecutive groups; trajectory classes: gains, loses, transient peak, stable; onset = first group exceeding the global average.

**Supplementary Table 2.** Sequences of pathway activation/deactivation along LIHC tumor progression (G4→G2→G1→G3). Representative pathways per mechanism, with average effective T frequency in each group, trajectory class, and onset.

| Mechanism | Representative pathway | G4 | G2 | G1 | G3 | Trajectory | Onset |
|---|---|---|---|---|---|---|---|
| Neurotrophic | NTF3 activates NTRK2 (TRKB) signaling | 0.83 | 0.87 | 0.91 | 0.93 | Gains | G2 |
| Senescence evasion | Evasion of Oncogene Induced Senescence Due to p14ARF Defects | 0.73 | 0.83 | 0.82 | 0.9 | Gains (plateau) | G2 |
| DNA synthesis | Plus-strand DNA synthesis | 0.64 | 0.75 | 0.82 | 0.96 | Gains | G1 |
| Glycosylation | Defective MOGS causes CDG-2b | 0.29 | 0.73 | 0.78 | 0.97 | Gains | G2 |
| Cell cycle (G2/M) | Phosphorylation of proteins in G2/M by Cyclin A:Cdc2 | 0.35 | 0.62 | 0.77 | 0.99 | Gains | G2 |
| Immune evasion | Blockage of phagosome acidification | 0.59 | 0.62 | 0.73 | 0.9 | Gains | G1 |
| DNA repair (MMR) | Defective Mismatch Repair Associated With MSH6 | 0.1 | 0.62 | 0.73 | 0.95 | Gains | G2 |
| Lipid metabolism (LXR) | NR1H2 & NR1H3 regulate gene expression linked to gluconeogenesis | 0.36 | 0.57 | 0.71 | 0.9 | Gains | G2 |
| Selenium metabolism | Methylation of MeSeH for excretion | 0.5 | 0.77 | 0.79 | 0.76 | Transient peak (G1) | G2 |
| Solute transport | Defective SLCO2B3 causes HBLRR | 0.67 | 0.69 | 0.64 | 0.76 | Transient peak (G3) | G4 |

**Interpretation.** The sequence is biologically coherent: neurotrophic signaling and senescence evasion are gained early (G2); DNA synthesis and immune evasion in G1; cell-cycle (G2/M), glycosylation and MMR in G2→G1; and the strongest activation of cell-cycle and MMR programs occurs in G3 (the most advanced group by active T-gene count). The selenium pathway shows a transient peak in G1, and solute transport peaks in G3.

#### S1.3 Level 3 — MST taxonomy and molecular subtypes of LIHC tumors

The MST constructed from the effective T frequency vectors of LIHC tumors produces a simpler structure than in PRAD. There are **two main branches**, plus three residual branches containing 2, 1, and 1 samples respectively:

- **Branch B1: 320 samples (~85% of the cohort)**, extending to approximately 12,000 activated T-genes.
- **Branch B2: 50 samples (~13% of the cohort)**, extending to approximately 8,000 activated T-genes.

The dominance of a single branch in LIHC contrasts with PRAD, where six branches contained 81% of the tumors. This suggests that LIHC is more homogeneous at the level of pathway deregulation, with a single dominant molecular trajectory and a minority alternative trajectory. As in PRAD, the extension of a branch in the temporal coordinate (number of activated T-genes) is correlated with its sample size, which is consistent with a sampling effect.

**Supplementary Table 3.** Main pathway deregulation programs in LIHC tumors. For each subtype: representative pathway, effective T frequency in B1 (320), in B2 (50), global frequency, and enrichment ratio (subtype/global).

| Program | Representative pathway | Freq. B1 (320) | Freq. B2 (50) | Freq. global | Enrichment (B1/global) | Enrichment (B2/global) |
|---|---|---|---|---|---|---|
| Neurotrophic | NTF3 activates NTRK2 (TRKB) signaling | 0.91 | 0.72 | 0.88 | 1.04 | 0.82 |
| Senescence evasion | Evasion of Oncogene Induced Senescence (p14ARF) | 0.82 | 0.7 | 0.8 | 1.02 | 0.88 |
| DNA synthesis | Plus-strand DNA synthesis | 0.82 | 0.76 | 0.8 | 1.02 | 0.95 |
| Glycosylation | Defective MOGS causes CDG-2b | 0.78 | 0.42 | 0.73 | 1.08 | 0.58 |
| Glycosylation | Defective ALG1 causes CDG-1k | 0.64 | 0.78 | 0.65 | 0.98 | 1.2 |
| Cell cycle (G2/M) | Phosphorylation of proteins in G2/M by Cyclin A:Cdc2 | 0.77 | — | 0.73 | 1.05 | — |
| Immune evasion | Blockage of phagosome acidification | 0.73 | 0.86 | 0.74 | 0.99 | 1.16 |
| DNA repair (MMR) | Defective Mismatch Repair Associated With MSH6 | 0.73 | — | 0.68 | 1.07 | — |
| Lipid metabolism (LXR) | NR1H2 & NR1H3 regulate gluconeogenesis | 0.71 | 0.37 | 0.66 | 1.08 | 0.56 |

| | | | | | | |
|---|---|---|---|---|---|---|
| Selenium metabolism | Methylation of MeSeH for excretion | 0.79 | 0.68 | 0.73 | 1.09 | 0.94 |
| Solute transport | Defective SLCO2A1 causes PHOAR2 | 0.76 | 0.39 | 0.39 | 1.97 | 1.01 |
| Metal sequestration | Metal sequestration by antimicrobial proteins | — | 0.58 | 0.42 | — | 1.38 |
| Invasion | Invadopodia formation | — | 0.47 | 0.38 | — | 1.25 |

**Interpretation.** The two main LIHC tumor subtypes show distinct functional profiles. The dominant subtype (B1) is characterized by neurotrophic signaling, evasion of senescence, and deregulation of lipid metabolism. The minority subtype (B2) is characterized by blockage of phagosome acidification, solute transport defects, and distinct glycosylation patterns.

#### S1.4 Relationship to the proliferative/non-proliferative dichotomy.

The two main branches identified by the MST in LIHC show a striking correspondence with the well-established proliferative/non-proliferative dichotomy of HCC [37]. The dominant branch (B1, 85% of tumors) is characterized by senescence evasion, lipid metabolism (LXR), and neurotrophic signaling (NTRK), features that align with the non-proliferative, differentiated class of HCC. The minority branch (B2, 13%) is characterized by phagosome acidification blockade, solute transport defects, and invadopodia formation, features that align with the proliferative, aggressive class. This correspondence suggests that the N/T framework captures the functional programs that underlie the genomic dichotomy, providing a pathway-level interpretation of the classical proliferative/non-proliferative classification.

#### S1.5 Level 1 (global). Protection in LIHC-N (normal)

The global analysis of the 50 LIHC normal tissues reveals a protection profile dominated by cellular maintenance programs, with RTK signaling protection confined to a distinct functional subset. Among the mechanisms with the highest global effective frequencies are DNA repair via BER (POLB-dependent long patch Base Excision Repair, 0.54; Base Excision Repair, 0.26), fatty acid metabolism (alpha-linolenic and linoleic acid metabolism, 0.46), protein folding (Association of TriC/CCT with target proteins, 0.52), and ciliary trafficking (BBSome-mediated cargo-targeting to cilium, 0.52). In contrast, protection of the RTK signaling network—comprising 76 pathways, including EGFR, FGFR1–4, NTRK2/3, MET, KIT, PDGF, and IGF1R—shows only a moderate global frequency (0.24), reflecting the fact that this protection is not uniformly distributed across normal samples but concentrated in a specific branch (A1EI, frequency 1.00; enrichment 4.17 relative to global; see S1.7). These global averages define the baseline protection landscape of normal liver tissue against which the loss of protection in the normal-to-tumor transition (S1.8) is assessed, and they indicate that LIHC-N prioritizes genomic integrity, lipid metabolism, and protein homeostasis, while RTK protection is restricted to a functional subgroup whose loss may be relevant to the NTRK activation observed in the dominant tumor subtype.

#### S1.6 Level 2 – Sequences of protection loss in LIHC normal tissue

Hierarchical clustering of the 50 LIHC normal samples on their effective N frequency vectors identifies three groups, ordered by the number of active N-genes as **G2** → **G3** → **G1**. This order goes from the samples with the most active N-genes (most intact protection) to those with the fewest (most advanced somatic evolution). Following the general criterion established in the Methods (Section 2.6), we compare the average effective N frequency of each pathway across these ordered groups. A pathway is considered active in a group if its group frequency exceeds the global cohort average (Step 1); an appreciable change requires an absolute difference of at least 0.10 between consecutive groups; and the trajectory classes are *loses*, *transient peak*, and *stable*. In normal tissue, somatic evolution is a process of **protection loss**, so only *loss* trajectories are interpreted as biologically meaningful progression.

**Supplementary Table 4. Sequences of protection programs along LIHC normal groups (G2 → G3 → G1).** Representative pathways are shown with their average effective N frequency in each group, their trajectory class, and their onset of loss (first group falling below the global average).

| Mechanism | Representative pathway | Global freq. | G2 (16) | G3 (14) | G1 (20) | Trajectory | Onset of loss |
|---|---|---|---|---|---|---|---|
| **DNA repair (BER)** | POLB-Dependent Long Patch Base Excision Repair | 0.54 | **0.78** | **0.67** | 0.25 | Loses | G3 |
| **Base Excision Repair** | Base Excision Repair | 0.26 | **0.43** | **0.43** | 0.11 | Loses | G1 |
| **Protein folding** | Association of TriC/CCT with target proteins | 0.52 | **0.94** | **0.67** | 0.05 | Loses | G3 |
| **Ciliary trafficking** | BBSome-mediated cargo-targeting to cilium | 0.52 | **0.94** | **0.67** | 0.05 | Loses | G3 |
| **Fatty acid metabolism** | alpha-linolenic and linoleic acid metabolism | 0.46 | **0.89** | **0.5** | 0.05 | Loses | G3 |
| **RTK signaling (network)** | 76 pathways (EGFR, FGFR1–4, NTRK2/3, MET, KIT, PDGF, IGF1R) | 0.24 | 0 | **1** | 0 | Sampling effect (G3) | — |

**Interpretation.**

The sequence G2 → G3 → G1 reveals a coherent hierarchy of protection loss for the cellular maintenance programs, and a sampling artifact for the RTK network.

First, the programs responsible for **cellular maintenance** display a clear **loss** trajectory. Protein folding (TriC/CCT), ciliary trafficking (BBSome), and fatty acid metabolism are maximally protected in G2 (0.89–0.94), partially maintained in G3 (0.50–0.67), and nearly absent in G1 (0.05). DNA repair (BER) follows the same pattern: POLB-dependent long patch BER is highest in G2 (0.78), remains elevated in G3 (0.67), and falls sharply in G1 (0.25). This indicates that G2 represents the subgroup with the most intact maintenance protection, corresponding to the **B1 branch** identified in the MST (S1.7), which is characterized by DNA repair, protein folding, and fatty acid metabolism.

Second, the **RTK signaling network** (76 pathways, including EGFR, FGFR1–4, NTRK2/3, MET, KIT, PDGF, and IGF1R) shows an apparent peak in G3 (frequency 1.00). Because somatic evolution is a process of loss, this cannot be interpreted as a biological gain of protection. Instead, it reflects a **sampling effect**: G3 is composed of a subset of samples (the B2-like branch of the MST, S1.7) in which RTK protection happens to be intact. With only 14 samples in G3, the group average is dominated by this subgroup and does not represent a genuine progression state. This is consistent with the global picture of S1.5, where RTK protection shows a moderate global frequency (0.24) precisely because it is concentrated in a minority of normal samples rather than uniformly maintained.

The ordered groups do not represent a smooth temporal trajectory; rather, they reflect the coexistence of these two functional subgroups in the normal cohort.

#### S1.7 Level 3 (MST). The MST on the LIHC normal samples.

The MST algorithm applied on the 50 LIHC normal samples produces **7 branches**, all small:

- **B1: 23 samples**, extending to ~300 activated N-genes.
- **B2: 12 samples**, extending to ~250 activated N-genes.
- Residual branches: 5, 4, 3, 2, 1 samples.

Normal LIHC tissue is divided into two functional groups: one with protection of cellular maintenance (branch B1: DNA repair, chromatid cohesion, fatty acid metabolism, protein folding, trafficking) and one with complete protection of RTK signaling (branch B2). The loss of RTK protection in normals may be related to the activation of NTRK (freq T 0.91) in the dominant tumor subtype.

**Supplementary Table 5.** Protection mechanisms in normal LIHC tissue. For each program: representative pathway, frequency in B1 (23), in B2 (12), global, and enrichment (branch/global).

| Program | Representative pathway | Freq. B1 (23) | Freq. B2 (12) | Freq. global | Enrichment (B1/global) |
|---|---|---|---|---|---|
| DNA repair (BER) | POLB-Dependent Long Patch Base Excision Repair | 0.78 | — | 0.54 | 1.45 |
| Base Excision Repair | Base Excision Repair | 0.39 | — | 0.26 | 1.5 |
| Fatty acid metabolism | alpha-linolenic and linoleic acid metabolism | 0.7 | — | 0.46 | 1.51 |
| Protein folding | Association of TriC/CCT with target proteins | 0.7 | — | 0.52 | 1.34 |
| Ciliary trafficking | BBSome-mediated cargo-targeting to cilium | 0.7 | — | 0.52 | 1.34 |
| RTK signaling (network) | 76 pathways (EGFR, FGFR1–4, NTRK2/3, MET, KIT, PDGF, IGF1R) | — | 1 | 0.24 | 4.17 |

#### S1.8 Connection between protection loss in normals and progression gain in tumors.

The comparison between protection mechanisms in normal LIHC tissue (S1.5–S1.7) and progression programs in tumors (S1.1–S1.3) reveals a coherent connection consistent with the N/T framework: the loss of N-gene activity in normal tissue precedes and predisposes to the gain of T-gene activity in the dominant tumor program. A direct branch-to-branch comparison between the normal and tumor MST taxonomies (S1.7 and S1.3, respectively) makes this correspondence explicit.

**Branch-to-branch correspondence.**

The comparison reveals a one-to-one mapping. **Normal B1 → Tumor B1:** the maintenance-protected background (BER, TriC/CCT, BBSome, fatty acid metabolism) is the one whose loss corresponds to the gain of the dominant tumor program (NTRK, senescence evasion, LXR).

**Normal B2 → Tumor B2:** the RTK-protected background (RTK network, 1.00) is the one whose loss corresponds to the gain of the minority tumor program (phagosome blockade, solute transport defects, invadopodia). This branch-level mapping is more informative than a simple global comparison, because it shows that the two tumor subtypes are not arbitrary partitions but the direct evolutionary continuation of the two normal protection profiles.

**Biological interpretation.** The dominance of tumor B1 (85%) over tumor B2 (13%) indicates that the loss of maintenance protection (normal B1) is the principal route to LIHC, whereas the loss of RTK protection (normal B2) defines a minority but biologically distinct trajectory. This dual correspondence is consistent with the proliferative/non-proliferative dichotomy of HCC [37].

The connection between protection loss and tumor progression is further supported by the metabolic reprogramming observed in the dominant subtype. The normal protection of fatty acid metabolism (alpha-linolenic/linoleic, 0.46) and protein folding (TriC/CCT, 0.52) is lost along the **G2 → G3 → G1** sequence, and tumor B1 acquires LXR-mediated lipid deregulation (0.71) and NTRK activation (0.91). These observations are consistent with metabolic classifications of HCC that identify glycolysis/cholesterol synthesis and N-glycan/mevalonate/sphingolipid programs as oncogenic axes [38], reinforcing the connection between protection loss in normals and metabolic reprogramming in the dominant tumor program.

---

### S2. Lung Adenocarcinoma (LUAD)

#### S2.1 Level 1 — Pathways important in LUAD tumors (global)

**Supplementary Table 6**. Global averages over the LUAD tumor cohort. Representative pathways:

| Mechanism | Representative pathway | Global freq. |
|---|---|---|
| Surfactant | Defective pro-SFTPC causes SMDP2 and RDS | 0.97 |
| Selenium metabolism | Methylation of MeSeH for excretion | 0.9 |
| Glycosylation | Defective ALG8 causes CDG-1h | 0.89 |
| Glycosylation | Reactions specific to the hybrid N-glycan synthesis pathway | 0.87 |
| FGFR signaling | FGFR4 mutant receptor activation | 0.84 |
| Glycosylation | Defective MOGS causes CDG-2b | 0.82 |
| Cell cycle | G2/M DNA replication checkpoint | 0.71 |
| DNA repair (HR) | Defective DNA double strand break response due to BRCA1 loss of function | 0.66 |

| | | |
|---|---|---|
| RTK | MET Receptor Activation | 0.51 |
| Lipid metabolism | LDL remodeling | 0.78 |

#### S2.2 Level 2 — Sequences of activation/deactivation in LUAD tumors

Hierarchical clustering of LUAD tumor samples identifies **four groups**, ordered by the number of active T-genes as **G2 → G3 → G1 → G4**. Comparing pathway averages across these ordered groups reveals a staggered gain of programs.

**Supplementary Table 7.** Sequences of pathway activation/deactivation along LUAD tumor progression (G2→G3→G1→G4).

| Mechanism | Representative pathway | G2 | G3 | G1 | G4 | Trajectory | Onset |
|---|---|---|---|---|---|---|---|
| Surfactant | Defective pro-SFTPC causes SMDP2 and RDS | 0.85 | 0.99 | 1 | 1 | Gains (plateau) | G2 |
| Selenium | Methylation of MeSeH for excretion | 0.61 | 0.98 | 0.99 | 0.9 | Gains (plateau) | G2 |
| Glycosylation (ALG8) | Defective ALG8 causes CDG-1h | 0.72 | 0.88 | 0.94 | 0.96 | Gains | G2 |
| Glycosylation (MOGS) | Defective MOGS causes CDG-2b | 0.73 | 0.8 | 0.91 | 0.7 | Transient peak (G1) | G2 |
| FGFR4 | FGFR4 mutant receptor activation | 0.68 | 0.89 | 0.86 | 0.91 | Gains | G2 |
| Cell cycle (G2/M) | G2/M DNA replication checkpoint | 0.27 | 0.47 | 0.94 | 0.8 | Gains (peak G1) | G3 |
| DNA repair (HR) | Defective DNA double strand break response due to BRCA1 | 0.34 | 0.9 | 0.82 | 0.9 | Gains (plateau) | G3 |
| RTK (MET) | MET Receptor Activation | 0.34 | 0.55 | 0.59 | 0.6 | Gains | G2 |
| Lipid | LDL remodeling | 0.65 | 0.85 | 0.84 | 0.66 | Transient peak (G3–G1) | G2 |

**Interpretation.** The sequence shows early gain (G2) of surfactant, selenium, glycosylation and FGFR4 programs; a strong peak of cell-cycle (G2/M) and HR repair in G1; and a late plateau (G4) for most programs. The transient behavior of MOGS and LDL remodeling reflects heterogeneity within the cohort rather than a monotonic progression.

#### S2.3 Level 3 — MST taxonomy in LUAD (single dominant group)

The MST constructed from the effective T frequency vectors of LUAD tumors **does not produce multiple main branches**, in contrast to PRAD (six main branches) and LIHC (two main branches). Instead, the LUAD cohort is dominated by **a single main branch** (plus 2 isolated samples), sharing a common core of deregulated programs. This lack of clear substructure is not a limitation of the method, but a biological property of LUAD: the tumor converges toward a common progression program that includes surfactant defects (freq 0.97), extensive glycosylation defects (0.53–0.89), selenium metabolism (0.91), FGFR1–4 signaling (0.50–0.84), and intense cell-cycle deregulation

(0.40–0.71). This convergence is consistent with the observation that the number of genes in the complete panel decreases progressively from PRAD to LIHC to LUAD [3] (and further to GBM), suggesting that the molecular heterogeneity of the tumor decreases as the tumor becomes more aggressive and evolved.

#### S2.4 Level 1 (global). Protection in LUAD-N (normal)

The global analysis of the 59 normal LUAD tissues reveals a protection profile dominated by stress-response, growth-signaling, and structural programs, with a notable absence of uniformly high protection across the cohort. Among the mechanisms with the highest global effective frequencies are TNFR1 signaling (TNF signaling, 0.58), autophagy and hypoxia (Cellular response to hypoxia and Macroautophagy, both 0.54), PTK6 signaling (PTK6 Expression, 0.54), surfactant (Defective pro-SFTPC causes SMDP2 and RDS, 0.44), and N-glycosylation (Sialic acid metabolism, 0.34). In contrast, cell cycle regulation (Mitotic Prophase, 0.27), NOTCH signaling (NOTCH1/4 Intracellular Domain Regulates Transcription, 0.22), PRC2/E2F6 (PRC2 methylates histones and DNA, 0.22), and FGFR2 signaling (FGFR2 mutant receptor activation, 0.19) show only moderate global frequencies. Notably, the MAPK/RAS/RAF axis (Oncogenic MAPK signaling, 0.31) and FGFR2 (0.19) are present but not dominant at the global level, reflecting the fact that these growth-signaling protections are concentrated in a specific branch.

These global averages define the baseline protection landscape of normal lung tissue against which the loss of protection in the normal-to-tumor transition (S2.5) is assessed, and they indicate that LUAD-N prioritizes stress/survival responses and autophagy/hypoxia programs, while growth-signaling protection (MAPK/FGFR2) and structural/developmental protection (surfactant/NOTCH) are restricted to specific functional subgroups whose loss may be relevant to the tumor programs observed in LUAD.

#### S2.5 Level 2. Hierarchical clustering of LUAD normal samples

Hierarchical clustering of the 59 LUAD normal samples on their effective N frequency vectors identifies two groups, ordered by the number of active N-genes as **G2 → G1**.

**Supplementary Table 8. Sequences of protection programs along LUAD normal groups (G2 → G1).** Representative pathways from the global analysis (S2.4) are shown with their average effective N frequency in each group, their trajectory class, and their onset of loss (first group falling below the global average).

| Mechanism | Representative pathway | Global freq. | G2 (18) | G1 (41) | Trajectory | Onset of loss |
|---|---|---|---|---|---|---|
| **TNFR1 signaling** | TNF signaling | 0.58 | **0.72** | 0.51 | Loses | G1 |
| **Autophagy** | Macroautophagy | 0.54 | **0.78** | 0.44 | Loses | G1 |
| **Hypoxia (HIF)** | Cellular response to hypoxia | 0.54 | **0.78** | 0.44 | Loses | G1 |
| **PTK6** | PTK6 Expression | 0.54 | **0.78** | 0.44 | Loses | G1 |
| **Cell cycle** | Mitotic Prophase | 0.27 | **0.83** | 0.02 | Loses | G1 |
| **Nucleosome assembly** | Nucleosome assembly | 0.54 | **0.94** | 0.37 | Loses | G1 |
| **Surfactant** | Defective pro-SFTPC causes SMDP2 and RDS | 0.44 | **0.5** | 0.41 | Loses (plateau) | G1 |
| **MAPK/RAS/** | Oncogenic MAPK signaling | 0.31 | **0.5** | 0.22 | Loses | G1 |

| | | | | | | |
|---|---|---|---|---|---|---|
| **RAF** | | | | | | |
| **FGFR2** | FGFR2 mutant receptor activation | 0.19 | **0.28** | 0.12 | Loses | G1 |
| **NOTCH** | NOTCH1/4 Intracellular Domain Regulates Transcription | 0.22 | **0.28** | 0.2 | Loses | G1 |
| **FGFR1** | Signaling by FGFR1 | 0.31 | 0.22 | **0.34** | Transient peak (G1) | — |

**Interpretation.**
The sequence G2 → G1 reveals a coherent hierarchy of protection loss for the LUAD normal compartment.

First, the programs of **stress/survival** (TNFR1 signaling, autophagy, hypoxia, PTK6) and **growth signaling** (MAPK/RAS/RAF, FGFR2) are maximally protected in G2 (0.28–0.78) and are lost in G1 (0.12–0.51). The most dramatic losses are observed in cell cycle (Mitotic Prophase: G2 0.83 → G1 0.02) and nucleosome assembly (G2 0.94 → G1 0.37), indicating that G1 represents a state of advanced somatic evolution in which cell-cycle and chromatin protection have been largely eroded.

Second, **surfactant** (structural/developmental) also shows a loss trajectory (G2 0.50 → G1 0.41), although with a smaller magnitude. This is consistent with the global profile of S2.4, where surfactant is moderately protected (0.44) and its loss precedes its strong deregulation in tumors (0.97; S2.1).

Third, **FGFR1** shows a *transient peak* in G1 (0.34), which is higher than its global average (0.31) and its G2 value (0.22). This is interpreted as a **sampling effect**: G1 is enriched for a subset of samples (the B3-like growth signaling branch of S2.6) in which FGFR1 protection happens to be intact, rather than as a biological gain. This is consistent with the global picture of S2.4, where FGFR1 is not a dominant protection program (global 0.31) and is concentrated in a specific branch.

The group-level sequence explains the global averages of S2.4: the moderate global frequencies of TNFR1 (0.58), autophagy/hypoxia (0.54), and PTK6 (0.54) are the result of averaging a higher frequency in G2 (0.72–0.78) and a lower frequency in G1 (0.44–0.51). This confirms that the normal LUAD compartment is functionally divided into two subgroups: one that maintains stress/survival and growth signaling protection and one that has lost these protections but maintains structural/developmental programs such as surfactant and FGFR1. The ordered groups do not represent a smooth temporal trajectory; rather, they reflect the coexistence of these functional subgroups in the normal cohort.

**Caveat on group balance.** The groups are unbalanced (G2 = 18, G1 = 41 samples). The hierarchy of protection in LUAD is supported primarily by the global averages of S2.4 while the group-level sequence is reported as a qualitative, complementary view.

#### S2.6 Level 3. The MST on LUAD normal samples.

Application of the MST algorithm to the 59 LUAD normal samples produces **three main branches** of comparable size, all extending to ~ 160 activated N-genes:

- **B1: 15 samples —** "structural/developmental" profile.
- **B2: 20 samples —** "stress/survival" profile.

- **B3: 14 samples** — "growth signaling" profile.
- 6 minor branches: 10 samples in total.

**Supplementary Table 9. Protection mechanisms in normal LUAD tissue.** For each program: representative pathway, frequency in B1 (15), B2 (20), B3 (14), global frequency, and enrichment ratio (best branch/global).

| Program | Representative pathway | Freq. B1 (20) | Freq. B2 (15) | Freq. B3 (14) | Freq. global | Enrichment (best branch/global) |
|---|---|---|---|---|---|---|
| Surfactant | Defective pro-SFTPC causes SMDP2 and RDS | 0.41 | 0.73 | 0.36 | 0.44 | 1.66 (B2) |
| NOTCH | NOTCH1/4 Intracellular Domain Regulates Transcription | 0.2 | 0.53 | 0.21 | 0.22 | 2.42 (B2) |
| PRC2/E2F6 | PRC2 methylates histones and DNA | 0.1 | 0.6 | 0.21 | 0.22 | 2.73 (B2) |
| Cell cycle | Mitotic Prophase | 0.18 | 0.47 | 0.36 | 0.27 | 1.72 (B2) |
| TNFR1 signaling | TNF signaling | 0.51 | 0.4 | 0.36 | 0.58 | — |
| Autophagy | Macroautophagy | 0.44 | 0.4 | 0.86 | 0.54 | 1.58 (B3) |
| Hypoxia (HIF) | Cellular response to hypoxia | 0.44 | 0.4 | 0.86 | 0.54 | 1.58 (B3) |
| N-glycosylation | Sialic acid metabolism | 0.27 | 0.4 | 0.36 | 0.34 | 1.18 (B2) |
| PTK6 | PTK6 Expression | 0.44 | 0.4 | 0.86 | 0.54 | 1.58 (B3) |
| MAPK/RAS/RAF | Oncogenic MAPK signaling | 0.22 | 0.13 | 0.71 | 0.31 | 2.34 (B3) |
| FGFR2 | FGFR2 mutant receptor activation | 0.11 | 0 | 0.57 | 0.19 | 4.21 (B3) |

Unlike LIHC, where the normal MST collapses into two main branches (one maintenance-protected and one RTK-protected), the LUAD normal compartment is partitioned into three functionally distinct branches. This tripartite structure indicates that normal lung tissue maintains three alternative protection profiles, which differ in the biological programs they prioritize.

**Interpretation.**
The three branches reveal a clear functional division of the normal LUAD compartment, which maps directly onto the group-level sequence of S2.5 and the global averages of S2.4.

**B1 – structural/developmental profile (15 samples).** B1 is characterized by the highest protection of TNFR1 signaling (0.51) and moderate protection of autophagy, hypoxia, and PTK6 (0.44 each), together with moderate surfactant (0.41) and cell cycle (0.18). This profile aligns with the branch that maintains a basal structural and stress-response state, without a dominant proliferative or growth-signaling program. In the group-level analysis (S2.5), B1 corresponds to the G1 subgroup, which has lost most of the stress/survival and growth signaling protections but retains surfactant and FGFR1.

**B2 – stress/survival profile (20 samples).** B2 is characterized by the highest protection of surfactant (0.73), NOTCH (0.53), PRC2/E2F6 (0.60), cell cycle (0.47), and N-glycosylation (0.40).

This profile aligns with a branch that prioritizes epithelial differentiation (surfactant), developmental signaling (NOTCH), chromatin regulation (PRC2), and a moderate proliferative capacity. In the group-level analysis (S2.5), B2 is the dominant component of the G2 subgroup, which shows high protection of stress/survival and cell-cycle programs.

**B3 – growth signaling profile (14 samples).** B3 is characterized by the highest protection of autophagy (0.86), hypoxia (0.86), PTK6 (0.86), MAPK/RAS/RAF (0.71), and FGFR2 (0.57). This profile aligns with a branch that prioritizes growth factor signaling (FGFR2/MAPK), metabolic adaptation to hypoxia (autophagy, HIF), and survival signaling (PTK6). In the group-level analysis (S2.5), B3 is the other major component of the G2 subgroup, and its high MAPK/FGFR2 protection explains the moderate global frequencies of these programs in S2.4 (0.31 and 0.19, with enrichment 2.34 and 4.21 relative to global).

**Biological significance.** The tripartite structure of the normal LUAD compartment is biologically meaningful because it identifies three distinct pre-tumoral protection profiles. B1 (structural/developmental) maintains a basal epithelial state; B2 (stress/survival) prioritizes surfactant, NOTCH, and chromatin regulation; and B3 (growth signaling) prioritizes FGFR2/MAPK, autophagy, and hypoxia adaptation. These three profiles are the normal baselines whose **loss** predisposes to the three TCGA transcriptional subtypes of LUAD (bronchioid, squamousid, and magnoid), as discussed in S2.7. The branch-level analysis therefore provides the missing link between the normal protection landscape and the tumor taxonomy.

#### S2.7 Comparison with existing LUAD classifications

The programs identified in our analysis show substantial correspondence with the TCGA transcriptional classification of LUAD [10]. The TCGA defines three transcriptional subtypes: **bronchioid**, **squamousid**, and **magnoid**, which differ in their dominant biological programs and clinical outcomes. In our framework, each TCGA subtype is interpreted as the tumor outcome of the **loss** of a specific normal protection profile, not as the direct continuation of that protection.

**Bronchioid subtype.** The bronchioid subtype is characterized by a structural/developmental profile, with low proliferation and expression of surfactant and NOTCH-related genes. In our normal tissue analysis (S2.5), branch **B1** (15 samples) is defined by a "structural/developmental" profile, with high protection of surfactant (0.41), NOTCH (0.20), and cell cycle (0.18). The **loss** of this structural/developmental protection may correspond to the bronchioid tumor, in which surfactant (0.97) and NOTCH are deregulated.

**Squamousid subtype.** The squamousid subtype is enriched for immune and stress-response signatures. In our normal tissue analysis, branch **B2** (20 samples) is defined by a "stress/survival" profile, with high protection of TNFR1 signaling (0.51) and autophagy (0.44). The **loss** of this stress-response protection may be related to the squamousid tumor, in which autophagy and hypoxia programs become moderately deregulated.

**Magnoid subtype.** The magnoid subtype is the most aggressive, characterized by high proliferation, RTK signaling, and genomic instability. In our normal tissue analysis, branch **B3** (14 samples) is defined by a "growth signaling" profile, with high protection of MAPK/RAS/RAF (0.71), FGFR2 (0.57), autophagy (0.86), and hypoxia (0.86). The **loss** of this growth signaling protection may correspond to the magnoid tumor, in which FGFR1–4 (freq T 0.50–0.84), MAPK/RAS/RAF (0.30–0.84), and cell cycle (0.40–0.71) are deregulated.

**Reconciling the single N/T group with the TCGA subtypes.** The TCGA classification subdivides LUAD into three transcriptional subtypes, whereas the N/T framework identifies a single dominant functional trajectory. These two observations are not contradictory: they reflect different levels of resolution. The TCGA subtypes capture qualitative differences in the dominant biological program (structural/developmental, stress/immune, or growth signaling), whereas the MST captures the qualitative convergence of LUAD toward a common progression program. The three protection profiles we identify in normal tissue (B1, B2, B3) represent distinct pre-tumoral baselines whose **loss** predisposes to the three TCGA subtypes, adding a temporal dimension to the static TCGA classification.

##### S2.8 Connection between protection loss in normals and progression gain in tumors

The comparison between protection in normals and progression in tumors of LUAD reveals several connections. The protection of surfactant is high in branch B2 (0.73) but the surfactant program is strongly deregulated in tumors (freq T 0.97), suggesting that loss of surfactant protection precedes its deregulation. The protection of MAPK/RAS/RAF and FGFR2 is high in branch B3 (0.71 and 0.57) but these programs are deregulated in tumors (freq T 0.30–0.84), suggesting that branch B3 may give rise to tumors with FGFR2/MAPK activation. Autophagy and hypoxia protection are high in branch B3 (0.86), but autophagy and hypoxia programs are moderately deregulated in tumors. The loss of NOTCH protection (branch B2) may contribute to the moderate NOTCH deregulation observed in tumors.

---

#### S3. Glioblastoma (GBM)

The TCGA GBM cohort contains very few normal samples, which limits the identification of N-genes and the analysis of protection. GBM tumors are therefore analyzed qualitatively, focusing on the main progression and protection programs.

##### S3.1 Level 1 — Pathways important in GBM tumors (global)

**Supplementary Table 10**. **Global averages over the GBM tumor cohort.** Main mechanisms with representative pathways and gobal frequencies.

| Mechanism | Representative pathway | Global freq. |
|---|---|---|
| TP53 loss | Loss of Function of TP53 in Cancer | 0.98 |
| Senescence evasion | Evasion of Oncogene Induced Senescence Due to p14ARF Defects | 0.4 |
| Cell cycle (G2/M) | G2/M DNA replication checkpoint | 0.98 |
| Lipid metabolism | Synthesis of CL (cardiolipin) | 1 |
| N-glycosylation | Defective ALG3 causes CDG-1d | 1 |
| N-glycosylation | Defective ALG8 causes CDG-1h | 0.99 |
| DNA repair (HR) | Impaired BRCA2 translocation to the nucleus | 0.99 |
| DNA repair (BER) | NEIL3-mediated resolution of ICLs | 0.99 |
| RTK | NTRK2 activates RAC1 | 0.9 |
| Apoptosis regulation | Activation of BMF and translocation to mitochondria | 0.94 |

| | | |
|---|---|---|
| Invasion | Invadopodia formation | 0.83 |
| ECM | Cell-extracellular matrix interactions | 0.82 |
| Selenium | Selenocysteine synthesis | 0.93 |

**Tissue-specific programs in GBM.** Several programs are notable for their high frequency and potential tissue specificity. The N-glycosylation defects (ALG3, ALG8, DPAGT1) are among the most universal features of the cohort (frequency > 0.99), suggesting that aberrant glycosylation is a core feature of GBM biology [39]. Selenium metabolism (selenocysteine synthesis, 0.93; selenoxamine acid metabolism, 0.90) is also highly activated, consistent with the role of selenium in redox homeostasis in the brain [40]. The deregulation of DNA repair pathways (BRCA2 0.99, NEIL3 0.99, Fanconi anemia 0.80) indicates that the normal repair program is replaced by an aberrant transcriptional program in tumors, which may contribute to therapeutic resistance.

#### S3.2 Levels 2 and 3. Sequences of pathways activation and taxonomy of GBM tumors.

For GBM, the hierarchical clustering groups of tumor samples leads to 7 groups that overlap and do not indicate a clear progression, and the MST collapses into a single branch that shares a common core of universally deregulated programs. This is consistent with the extreme aggressiveness and evolutionary stage of GBM. Therefore, **GBM-T is analyzed only at Level 1 (global)**.

The 7 groups of the hierarchical clustering may be seen as a sub-structure of the single MST branch.

#### S3.3 Correspondence with existing GBM classifications.

**Correspondence with the TCGA (Verhaak) classification.** The TCGA classification of GBM, established by Verhaak et al. [42], defines four transcriptional subtypes: classical, proneural, neural, and mesenchymal. The classical subtype is defined by EGFR amplification and activation, the proneural subtype by PDGFRA alterations and IDH1 mutations, and the mesenchymal subtype by NF1 loss and mesenchymal markers. Our data show prominent RTK signaling (NTRK2, FLT3, PDGFR) that cuts across these subtypes, reflecting the convergent dependence of GBM on RTK pathways. The near-universal TP53 loss we observe (frequency 0.97) is consistent with the high frequency of TP53 mutations across all GBM subtypes [42]. The lipid metabolism signature we detect (cardiolipin, sphingolipids) is particularly interesting in light of the proneural subtype's association with IDH1 mutations, which affect lipid metabolism.

**Reevaluation of the Neural subtype and the three-subtype framework.** The Neural subtype of Verhaak et al. has been subsequently questioned because its expression signature largely reflects contamination with normal brain tissue rather than a genuine tumor-intrinsic program. Under the WHO 2021 classification of central nervous system tumors [41], GBM is defined exclusively as IDH-wildtype, and the prevailing framework is the three-subtype model (proneural, classical, and mesenchymal), which excludes the Neural subtype. This reevaluation is consistent with the convergent RTK and cell-cycle programs we observe, which cut across the three remaining subtypes rather than defining a separate Neural class.

Note that the TCGA data were generated and annotated under older classification schemes. A direct comparison with these recent frameworks is therefore not entirely fair, and the correspondence reported above should be interpreted with this limitation in mind.

#### S3.4 Protection mechanisms in normal GBM tissue

Although the TCGA GBM cohort contains only 5 normal samples, the aggregate analysis of their effective N frequencies provides a qualitative view of the protection mechanisms in normal brain tissue. Note that the extremely high frequencies of some protection programs could be an artifact related to the small number of samples.

**Supplementary Table 11.** Protection mechanisms in normal GBM tissue (average over 5 normal samples).

| Program | Representative pathway | Avg. N freq. (5 samples) | Status |
|---|---|---|---|
| DNA repair (HR) | Defective DNA double strand break response due to BRCA1 loss of function | 0.8 | Maintained |
| DNA repair (BER) | POLB-Dependent Long Patch Base Excision Repair | 0.3 | Partially lost |
| Apoptosis | Activation, translocation and oligomerization of BAX | 0.8 | Maintained |
| Senescence | Evasion of Oncogene Induced Senescence Due to p16INK4A Defects | 1 | Maintained (protection against senescence evasion) |
| Cell cycle | Drug-mediated inhibition of CDK4/CDK6 activity | 1 | Maintained |
| RTK signaling | FGFR1b ligand binding and activation | 1 | Maintained |
| RTK signaling | Activated NTRK2 signals through FYN | 0.8 | Maintained |
| Metabolism (fatty acid) | mitochondrial fatty acid beta-oxidation of saturated fatty acids | 0.8 | Maintained |
| Metabolism (lipid) | Synthesis of CL (cardiolipin) | 1 | Maintained |
| Glycosylation | Defective ALG3 causes CDG-1d | 1 | Maintained |
| Immune | Blockage of phagosome | 0.8 | Maintained |

**Biological interpretation.** The 5 normal GBM samples show a distinct protection profile. The most robustly maintained mechanisms are those that prevent oncogenic transformation: DNA repair (HR, 0.80), apoptosis (BAX, 0.80), senescence protection (p16INK4A, 1.00), and cell cycle inhibition (CDK4/6, 1.00). RTK signaling (FGFR1, NTRK2) and lipid metabolism (cardiolipin, fatty acid oxidation) are also strongly maintained. In contrast, hypoxia (HIF) protection is partially lost (0.51), and surfactant protection is essentially absent (0.20), which is expected for brain tissue. The partial loss of autophagy (pexophagy, 0.60) and BER (0.30) suggests that these mechanisms are beginning to decline in the normal population.

#### S3.5 Connection between protection loss and tumor progression in GBM

The comparison between protection mechanisms in normal tissue and progression programs in tumors reveals several connections. The loss of apoptosis protection (normal N freq 0.80) is reflected in the near-universal TP53 loss in tumors (0.97) and the dysregulation of apoptotic pathways (BMF 0.94). The loss of senescence protection (normal N freq 1.00 for p16INK4A) is reflected in the evasion of senescence observed in a subset of tumors (p14ARF 0.40). The maintenance of RTK protection in normals (FGFR1 1.00, NTRK2 0.80) contrasts with the high RTK **deregulation** in tumors (NTRK2 0.90, FLT3 0.89), suggesting that loss of RTK protection precedes RTK-driven progression. Similarly, the maintenance of HR protection in normals (0.80) contrasts with the strong deregulation of DNA repair pathways in tumors (BRCA2 0.99, NEIL3 0.99), indicating that the normal repair protection is lost and replaced by an aberrant transcriptional program. The partial loss of hypoxia protection in normals (0.51) may predispose to the hypoxic adaptation observed in GBM. Finally, the maintenance of lipid metabolism protection in normals (cardiolipin 1.00) is contrasted by the intense lipid metabolism in tumors (cardiolipin 1.00, sphingolipids 0.70), suggesting that this program is reprogrammed rather than lost.

These observations support the N/T framework in GBM: the loss of protection mechanisms (apoptosis, senescence, RTK signaling, DNA repair, hypoxia) in normal tissue precedes and predisposes to the gain of progression programs in tumors (TP53 loss, senescence evasion, RTK activation, DNA repair deregulation, metabolic reprogramming). The limited number of normal samples (5) means that these conclusions are qualitative, but they are consistent with the known biology of GBM.

---

### Supplementary Figure Captions

**Supplementary Figure 1.** Clustering of tumor samples according to the effective T frequency vectors in LIHC. Upper panel: Effective frequencies of pathways. Central panel: Dendrogram. Lower panel: Number of activated pathways vs. active T-genes.

**Supplementary Figure 2.** Taxonomy of LIHC tumors according to the MST algorithm. Upper panel: The MST diagram. Lower panel: Number of samples in the main branches.

**Supplementary Figure 3.** Clustering of normal samples according to the effective N frequency vectors in LIHC. Upper panel: Effective frequencies of pathways. Central panel: Dendrogram. Lower panel: Number of activated pathways vs. active N-genes.

**Supplementary Figure 4.** Taxonomy of LIHC normal samples according to the MST algorithm. Upper panel: The MST diagram. Lower panel: Number of samples in the main branches.

**Supplementary Figure 5.** Clustering of tumor samples according to the effective T frequency vectors in LUAD. Upper panel: Effective frequencies of pathways. Central panel: Dendrogram. Lower panel: Number of activated pathways vs. active T-genes.

**Supplementary Figure 6.** Taxonomy of LUAD tumors according to the MST algorithm. The MST diagram shows a single main branch.

**Supplementary Figure 7.** Clustering of normal samples according to the effective N frequency vectors in LUAD. Upper panel: Effective frequencies of pathways. Central panel: Dendrogram. Lower panel: Number of activated pathways vs. active N-genes.

**Supplementary Figure 8.** Taxonomy of LUAD normal samples according to the MST algorithm. Upper panel: The MST diagram. Lower panel: Number of samples in the main branches.

**Supplementary Figure 9.** Clustering of tumor samples according to the effective T frequency vectors in GBM. Upper panel: Effective frequencies of pathways. Central panel: Dendrogram. Lower panel: Number of activated pathways vs. active T-genes.

**Supplementary Figure 10.** Taxonomy of GBM tumors according to the MST algorithm. The MST diagram shows a single main branch.

---

---

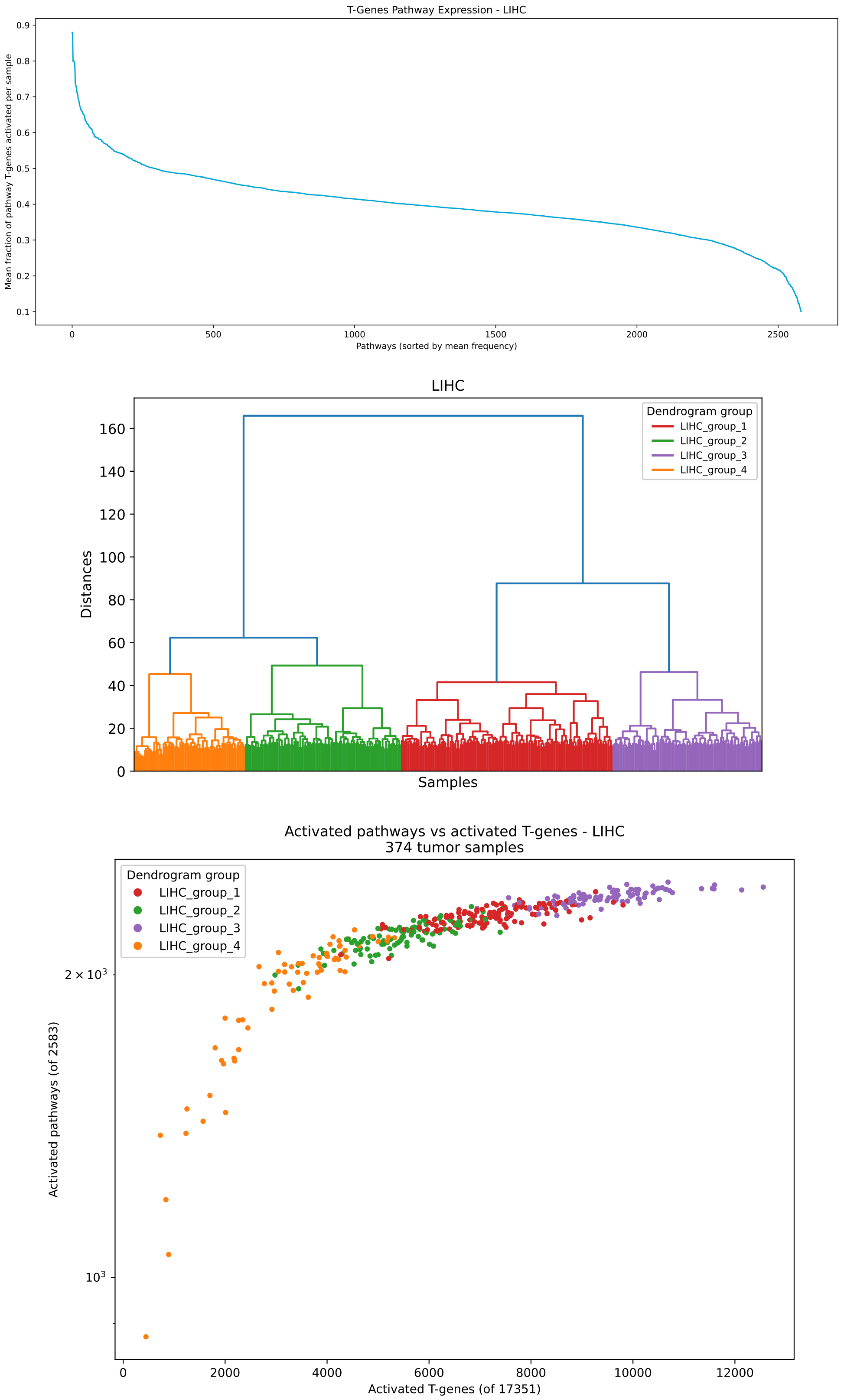


**Supplementary Figure 1.**

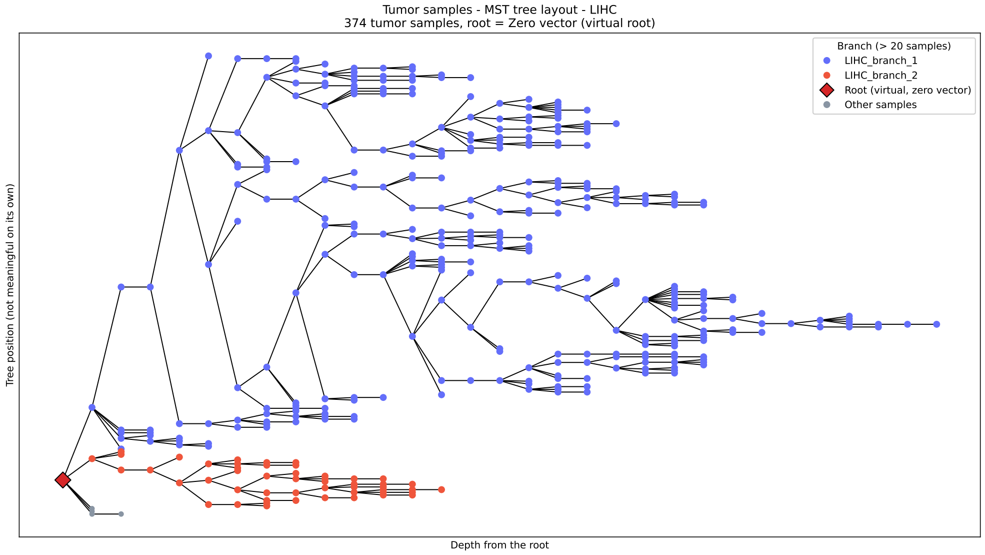


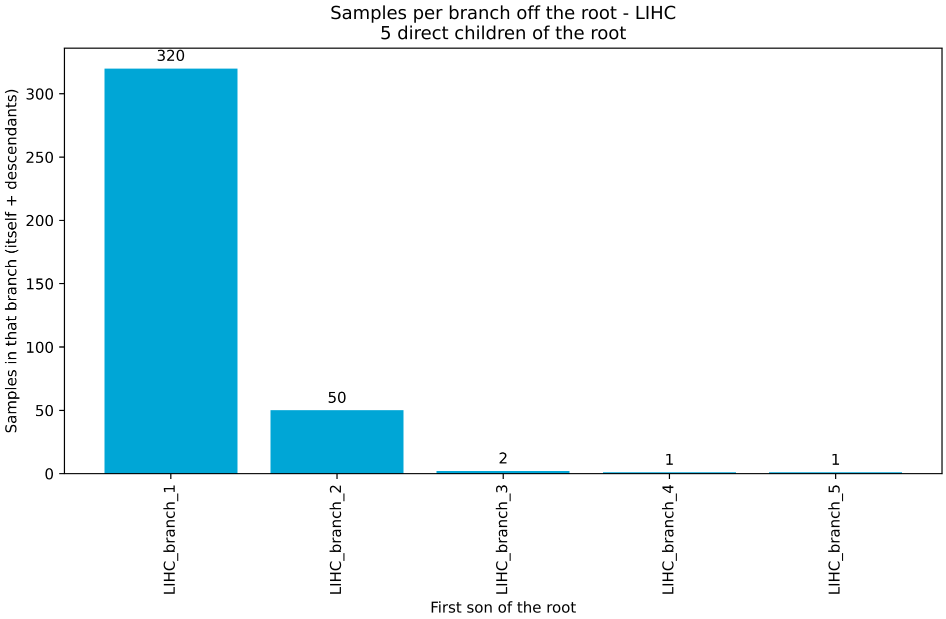


### Supplementary Figure 2.

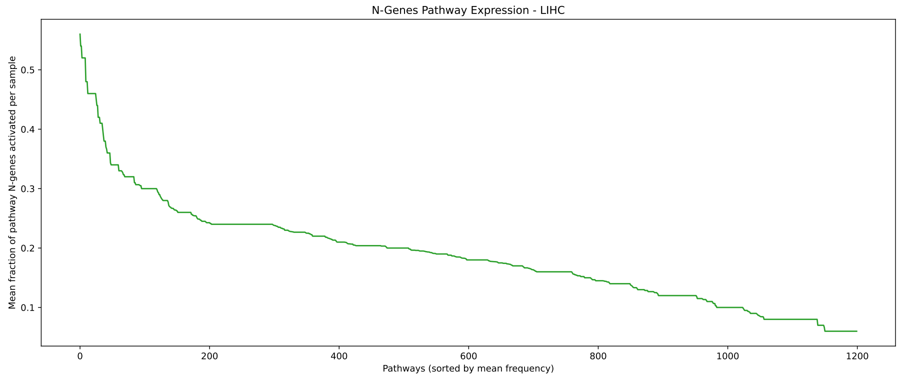


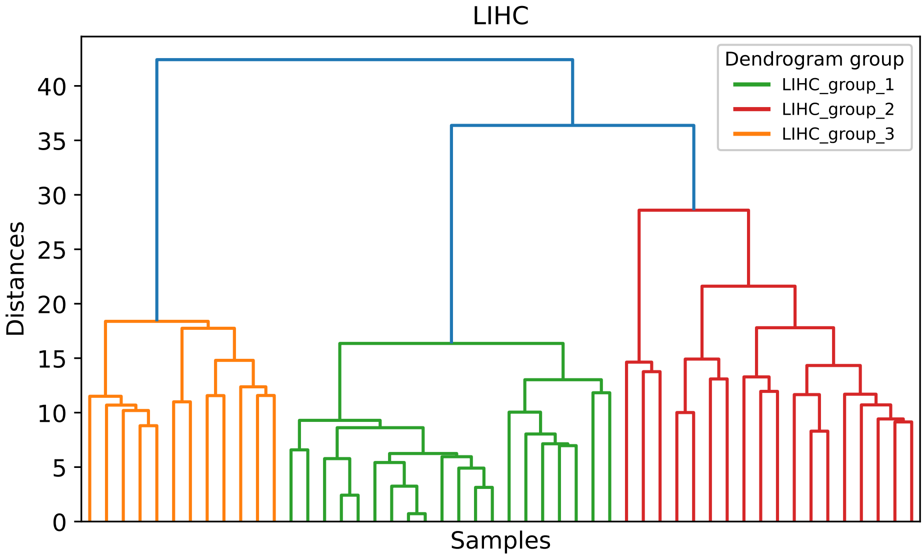


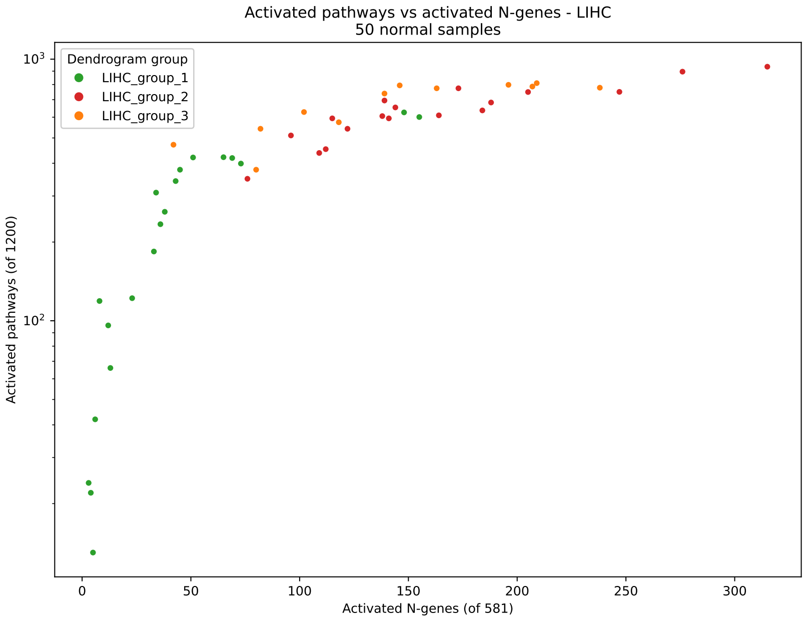


**Supplementary Figure 3.**

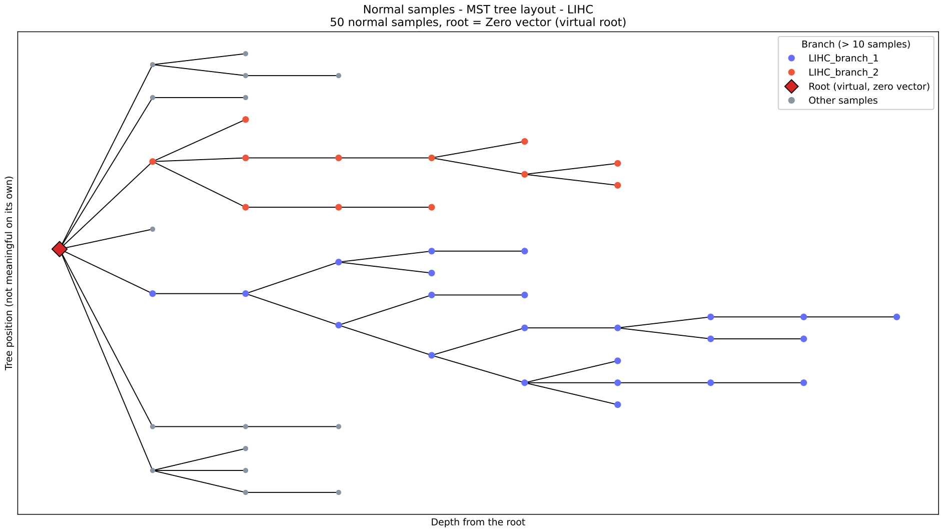


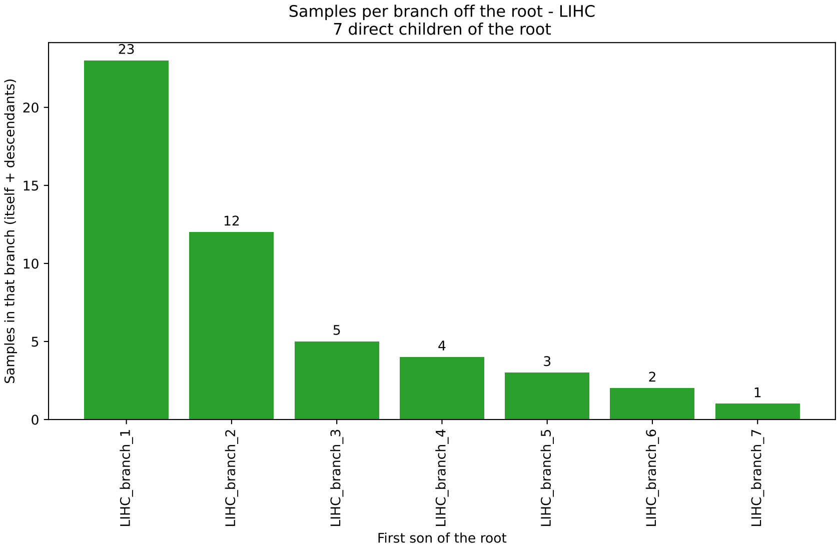


**Supplementary Figure 4.**

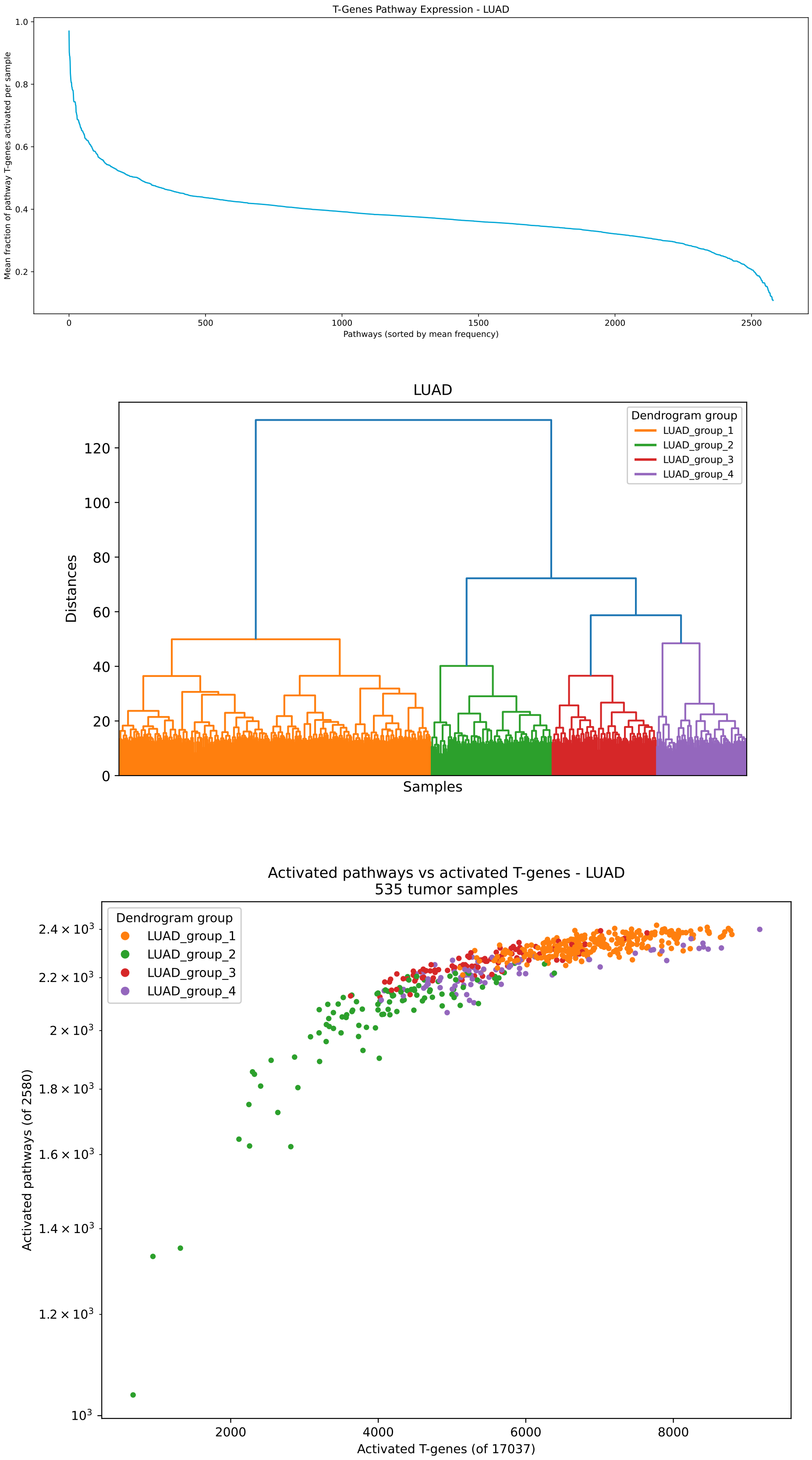


**Supplementary Figure 5.**

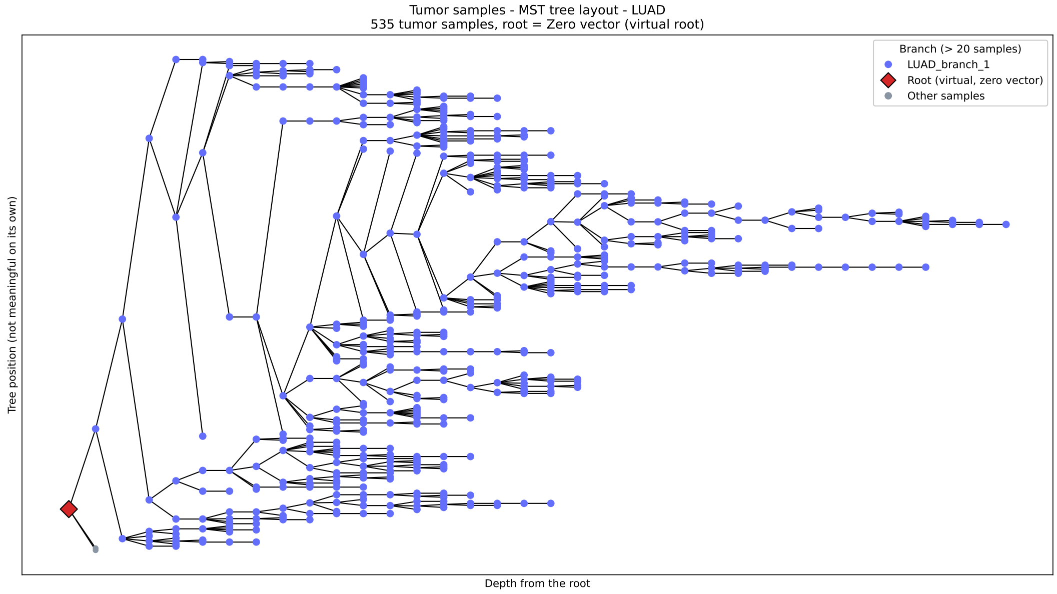


### Supplementary Figure 6.

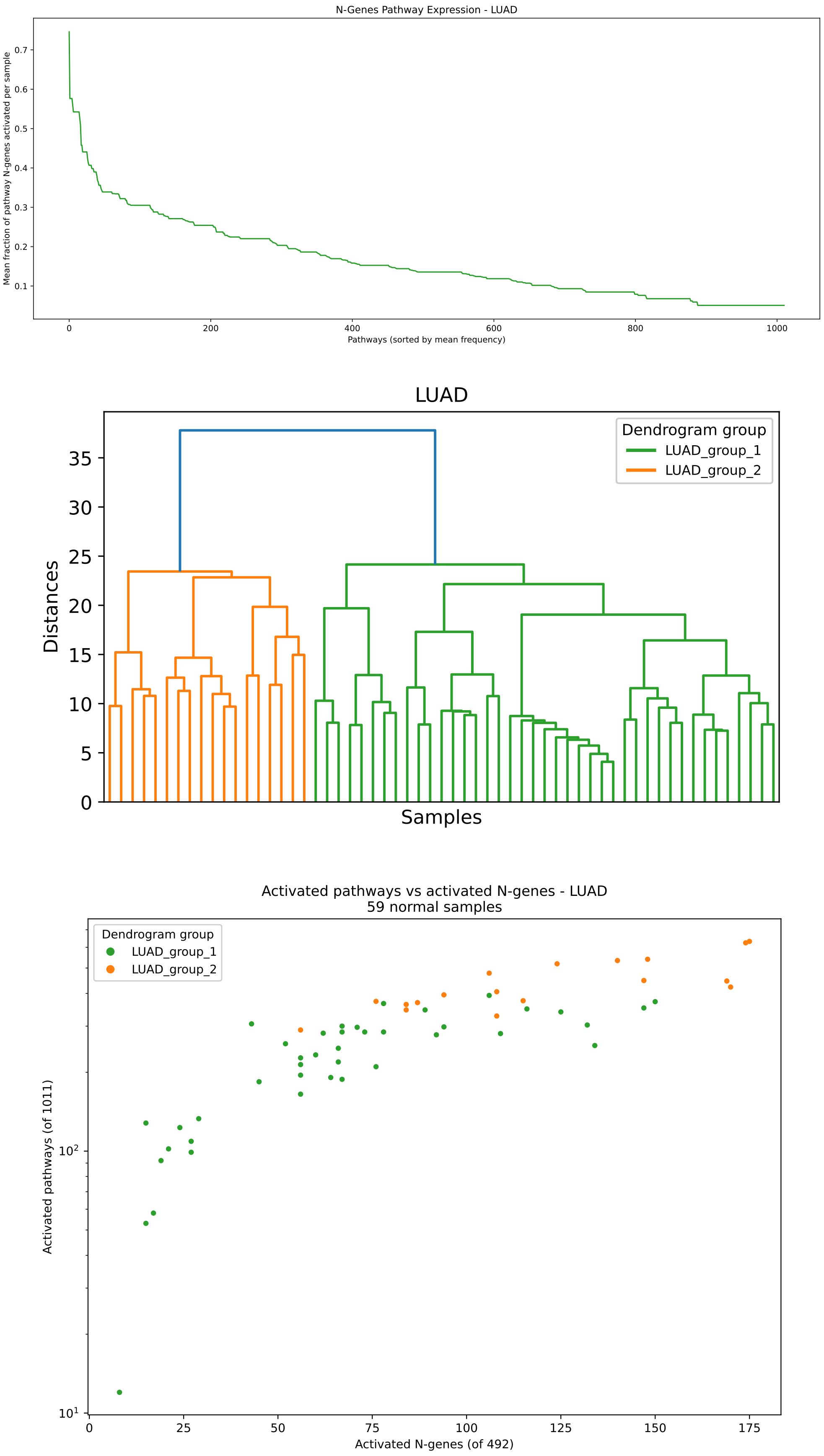


**Supplementary Figure 7.**

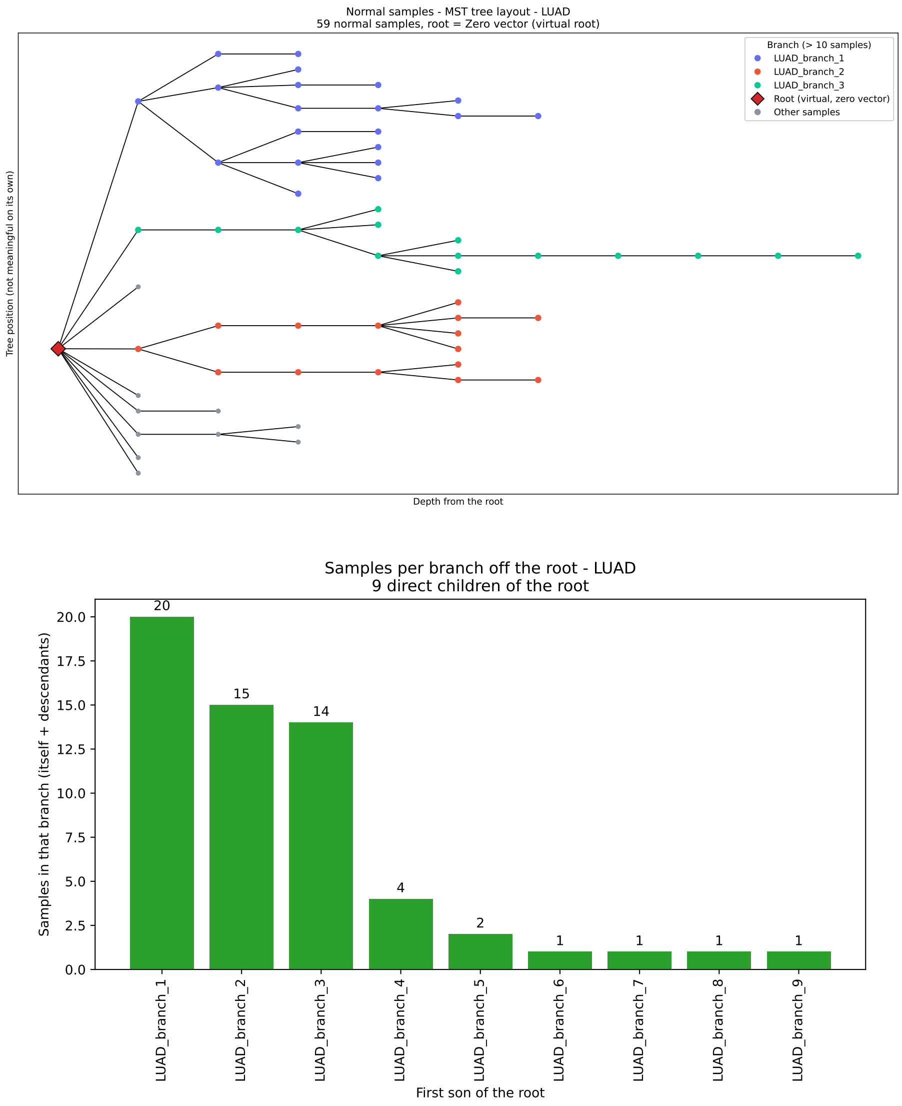


### Supplementary Figure 8.

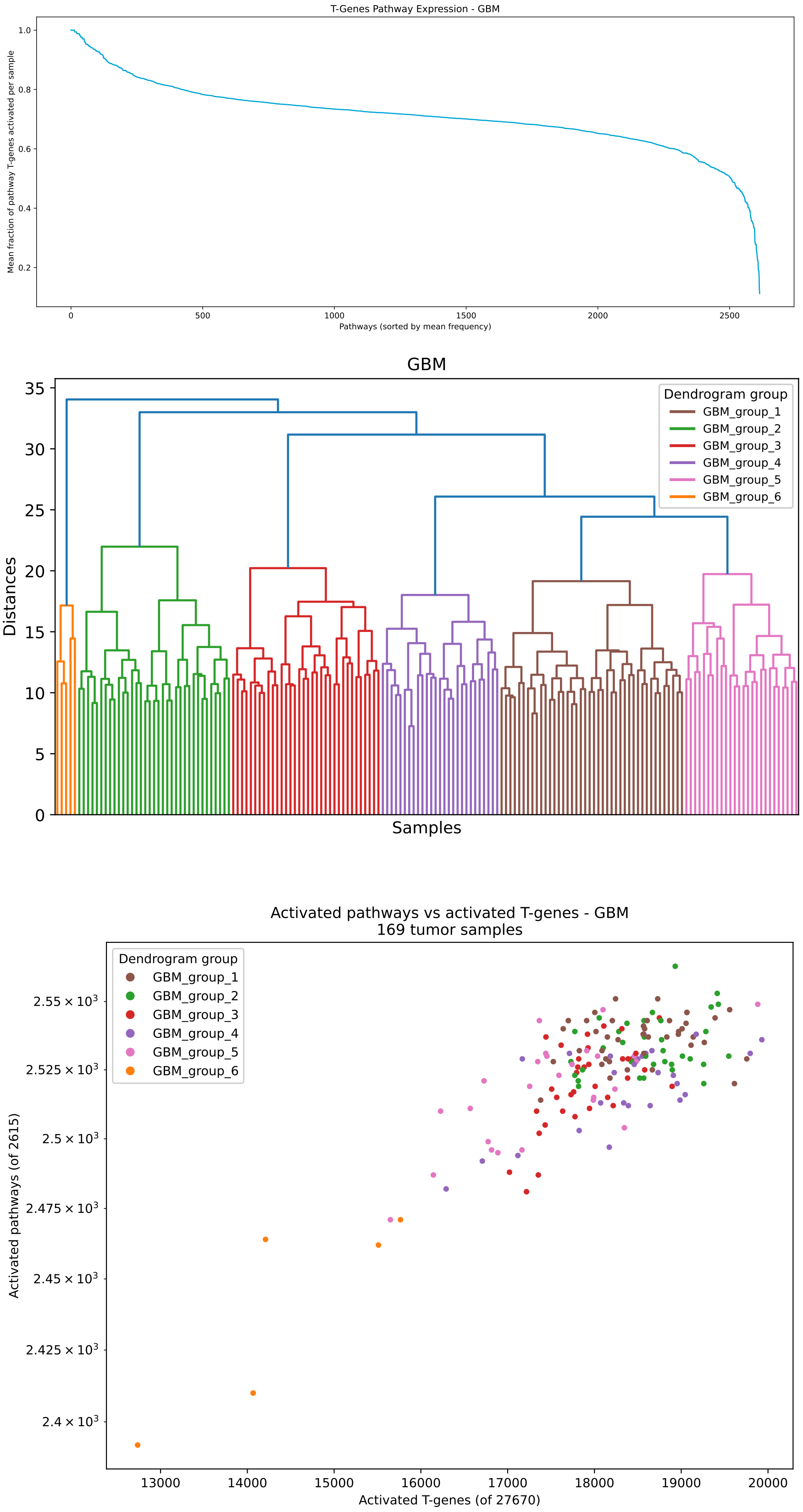


**Supplementary Figure 9.**

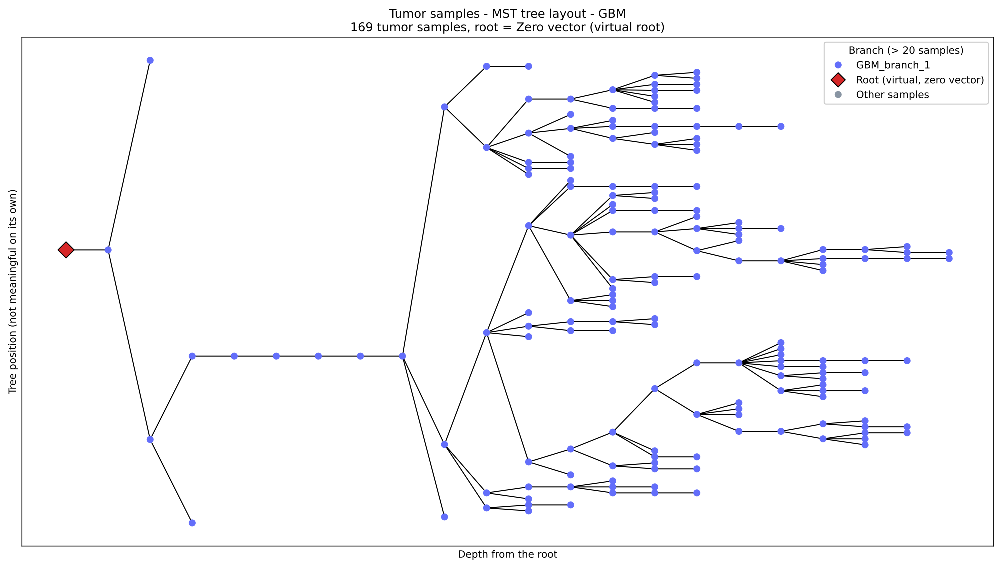


**Supplementary Figure 10.**
